# Detecting Genetic Interactions with Visible Neural Networks

**DOI:** 10.1101/2024.02.27.582086

**Authors:** Arno van Hilten, Federico Melograna, Bowen Fan, Wiro Niessen, Kristel van Steen, Gennady Roshchupkin

## Abstract

Non-linear interactions among single nucleotide polymorphisms (SNPs), genes, and pathways play an important role in human diseases, but identifying these interactions is a challenging task. Neural networks are state-of-the-art predictors in many domains due to their ability to analyze big data and model complex patterns, including non-linear interactions. In genetics, visible neural networks are gaining popularity as they provide insight into the most important SNPs, genes and pathways for prediction. Visible neural networks use prior knowledge (e.g. gene and pathway annotations) to define the connections between nodes in the network, making them sparse and interpretable. Currently, most of these networks provide measures for the importance of SNPs, genes, and pathways but lack details on the nature of the interactions. In this paper, we explore different methods to detect non-linear interactions with visible neural networks. We adapted and sped up existing methods, created a comprehensive benchmark with simulated data from GAMETES and EpiGEN, and demonstrated that these methods can extract multiple types of interactions from trained visible neural networks. Finally, we applied these methods to a genome-wide case-control study of inflammatory bowel disease and found high consistency of the epistasis pairs candidates between the interpretation methods. The follow-up association test on these candidate pairs identified seven significant epistasis pairs.

## Introduction

Machine learning methods, particularly neural networks, have been a disruptive technology that has transformed numerous fields in the last decade. Machine learning and deep learning have completely reshaped the fields of biomedical image segmentation (1), natural language processing (2, 3), protein folding (4) and many more. The rise of deep learning can be attributed to three main factors. First, sufficiently large neural networks can approximate any function (5, 6). Neural networks are thus not constrained to linear combinations, but can find and leverage non-linear interactions between inputs. Secondly, neural networks scale well with data set size (7). Neural networks thrive in large data sets with many examples as it allows the network to find complex patterns. Third, neural networks are flexible; their architecture can be easily modified for different tasks and different types of data. For the imaging domain, this led to convolutional neural networks (CNNs) while for natural language processing, transformers have deeply impacted the field (8).

In population-based genetics where there is a large number of input SNPs, there is a domain-specific trend to embed neural networks with prior biological knowledge, such as gene and pathway information, to create sparse and interpretable neural networks that predict genetic risk (9–13). These interpretable neural networks, coined visible neural networks (14), provided a solution to the two main challenges for neural networks for genetic data. The large number of input features - up to millions of SNPs - and the need for explainable methods. Prior knowledge such as gene and pathway information is embedded in the neural network architecture to define which node should connect and which not, resulting in a sparse and interpretable neural network. In these networks, each node represents a biological entity by the inputs it groups (e.g. SNPs are grouped by gene). The weights of the connections represent how predictive these entities (i.e. SNPs, genes and pathways) are for the final prediction. However, current methods for interpreting these networks, such as Layer-wise Relevance Propagation (15), Integrated Gradients (16) and DeepLIFT (17), only provide the importance for each entity and do not provide insight in the nature of the relation between entities. Attributing entities with an importance score, a single value for each input, provides an incomplete view of the decision process. Neural networks thrive because they can learn (non-linear) combinations of features and these cannot be expressed by a single importance value. Thus, for a more complete overview of the decision process of neural networks and to understand the nature of the relation between SNPs, genes, and pathways, it is important to detect and understand which input features interact with each other in neural networks.

Non-linear effects are ubiquitous in biology. Detecting and understanding these interactions is necessary to fully model the complex biological mechanisms that exist between genotype and phenotype (18, 19). Detecting interactions between genes and, in particular, SNPs (epistasis) comes with an inherent computational challenge. Following Fisher’s (20) definition of statistical epistasis, i.e., as a deviation from the additive expectation of allelic effects, the possible set of interactions exceeds *𝒪* (*p*^2^), with *p* the number of features. Thus, an exhaustive search is computationally unfeasible for a large number of inputs. In genetics, genome-wide association studies (GWASes) consider several millions of SNPs and detecting epistasis is thus infeasible to date without extra interventions to reduce the search space of possible interactions, due to the sheer number of possible combinations. Fortunately, there is a wide variety of non-exhaustive epistasis detection methods available. Epiblaster (21), takes a two-step approach that first searches for a smaller number of likely candidates using correlation before performing full rank logistic regression to confirm significance. MB-MDR (22), a non-parametric model often used to detect epistasis, conditions interaction testing on lower order effects. Other approaches, such as machine learning applications, build a prediction model first and then extract interaction information from this model. Tree-based classifiers can use the structure of the trees to find epistasis candidates (23, 24) or use the prediction model with permuted input data to find interacting features (25). For neural networks, the field of explainable AI (XAI) provides many tools that aim to explain the trained neural network. Most of these tools focus on input feature attribution and place little emphasis on finding interacting features. However, Greenside et al. (26) showed that feature attribution methods can be used to find epistasis candidates efficiently and Tsang et al. (27) proposed a method for finding statistically significant feature interactions using the weights of a neural network.

Combining these methods with visible neural networks might enhance our understanding of these neural networks and thereby the underlying biology. In this paper, we evaluate the performance and consistency of several post-hoc interpretation methods on visible neural networks from the GenNet framework (10). Primarily, we focus on epistasis detection, i.e., a pair of SNPs whose combination affects the phenotype. After training these networks, we apply: neural interaction detection (NID) (27), PathExplain (28) and Deep Feature Interaction Maps (DFIM) (26) to investigate how well these methods explain interactions learned by visible neural networks. Moreover, on the learned network, we analyze which gene gives a relative local improvement in predictive power (RLIPP) (13). We compare these methods to literature epistasis methods, such as light gradient-boosting machine, Epiblaster and MB-MDR, using simulated data from Epigen and Gametes. Finally, we apply these post-hoc interpretation methods to visible neural networks trained on data from the Inflammatory Bowel Disease Consortium.

## Materials

To evaluate epistasis methods in a controlled environment with known ground truth, we used simulated data from two different methods: GAMETES (29) for strict and pure epistasis models and EpiGEN (30) for more complex simulations. Finally, we applied the methods to the data from the International Inflammatory Bowel Disease Genetics Consortium (IIBDGC) to test the approaches in human data.

### GAMETES

is an open-source simulation package to generate pure and strict epistatic models, thus epistasis models without linkage disequilibrium and marginal effects but with two loci contributing to a discrete phenotype in a strictly non-linear manner. Fourteen different sets of simulations were simulated with varying sample sizes {3000, 12000}, heritability {0.05, 0.1, 0.2, 0.3}, and number of SNPs {25, 100, 1000}. An overview of the simulation settings can be found in Supplementary Table table 2.

**Table 1.** Properties of the different epistasis detection algorithms used and associated input. In detail, we describe the main category they belong to; if the test they perform is conditional on main effects; if they can naturally consider covariates, if the missing SNPs need to be imputed, and the type of epistasis score they yield.

| Method | Main category | Test conditional<br>on main effects | Covariates<br>adjustment | Imputation<br>required | Detect direct<br>interactions |
| --- | --- | --- | --- | --- | --- |
| Epiblaster (21) | Statistical | Yes | No | Yes | P-value |
| LightGBM (39) | ML | No | No | No | Interaction strength |
| MB-MDR (43) | Dimensionality reduction | Yes | Yes | No | P-value |
| NID (27) | ML, statistical | Yes | No | Yes | Interaction strength |
| DFIM (26) | ML | No | No | Yes | Interaction strength |
| PathExplain (28) | ML | No | No | Yes | Interaction strength |

**Table 2.** The thirteen simulation characteristics of the GAMETES simulations. Note that The first two simulations do not have interacting SNPs and that the phenotype is thus randomly defined.

| Simulation ID | Sample size | Input SNPs | Num. interacting SNPs | Num. simulations | Heritability |
| --- | --- | --- | --- | --- | --- |
| 1 | 3000 | 25 | 0 | 100 | 0.10 |
| 2 | 3000 | 25 | 0 | 100 | 0.20 |
| 3 | 3000 | 25 | 2 | 100 | 0.05 |
| 4 | 3000 | 25 | 2 | 100 | 0.10 |
| 5 | 3000 | 25 | 2 | 100 | 0.20 |
| 6 | 3000 | 50 | 2 | 100 | 0.10 |
| 7 | 3000 | 50 | 2 | 100 | 0.20 |
| 8 | 3000 | 100 | 2 | 100 | 0.10 |
| 9 | 3000 | 100 | 2 | 100 | 0.20 |
| 10 | 12000 | 100 | 2 | 10 | 0.10 |
| 11 | 12000 | 100 | 2 | 10 | 0.20 |
| 12 | 12000 | 1000 | 2 | 10 | 0.10 |
| 13 | 12000 | 1000 | 2 | 10 | 0.20 |
| 14 | 12000 | 1000 | 2 | 10 | 0.30 |

### EpiGEN

on the other hand, is a simulation pipeline built to simulate more complex phenotypes based on realistic genotype data. For example, EpiGEN allows the use of HAPGEN2 to simulate genotype data with similar characteristics (linkage disequilibrium, ethnicity, etc.). Additionally, EpiGEN was used to explore the effects of different epistasis models and SNPs with marginal effects. Using HAPGEN2 as a basis, we created simulations with varying sample sizes {3000, 12000}, number of SNPs {100, 1000}, interactions models {*joint*-*dominant, joint*-*recessive, multiplicative* and *exponential}*, and interaction strength {3, 10, 100}. Different interaction models mimic different structures of the epistasis. An interaction strength of, e.g., 10, means that an individual with the epistatic pair has 10-times the risk of someone without. Overall, we generated 384 different simulations: 288 with a marginal background effect and 96 pure epistasis models where only interaction effects lead to the response. All simulation parameters for EpiGEN can be found in Supplementary Table table 3. For in-depth details on the simulations, we refer to the original paper (30).

**Table 3.**
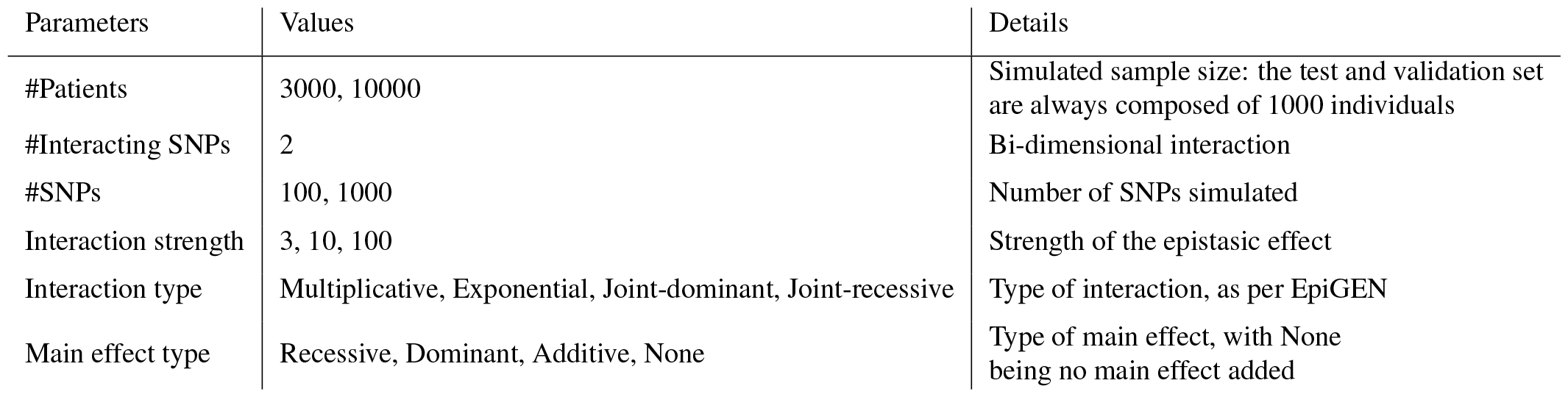
Hyperparameters used in the EpiGEN simulations.

### IBD dataset

We investigated the IBD dataset from the International Inflammatory Bowel Disease Genetics Consortium (IIBDGC). The data contains cases with non-infectious inflammations of the bowel, including Ulcerative colitis (UC) and Crohn’s disease (CD), the two main categories of IBD (31). The dataset was genotyped on the Immunochip SNP array (32). We performed quality control as in Ellinghaus, et al. (33), reducing the number of SNPs from 196 524 to 130 071. The final dataset contained 66 280 samples, including 32 622 cases (individuals with IBD) and 33 658 controls. Since the IIBDGC dataset aggregates multiple cohorts, confounders by shared genetic ancestry is a concern. As in Ellinghaus et al. (33), we used the first 7 principal components to model population stratification. We adjusted the phenotypes for epistasis detection methods that cannot include covariates. The same quality control steps as in Duroux, et al. (34) were applied. We removed rare variants (MAF *<* 5%) or in Hardy–Weinberg equilibrium (p-value *<* 0.001). All risk SNPs described in Liu et al. (35) were included.

## Methods

### Visible neural networks

The GenNet framework (10) was used to create sparse and interpretable neural networks. These visible neural networks use biological knowledge embedded in the neural network architecture to define connections between nodes. Figure 1 illustrates the employed neural network architecture. Each network had a depth of three or four layers, depending on the input encoding, and was structured according to Figure 1. Several changes were made to the original framework to improve the neural networks for epistasis detection. First, we tested one-hot input encoding in addition to the standard (additive) input encoding. Secondly, we added multiple filters for each gene to allow the network to find and use multiple patterns per SNP and gene, followed by an extra layer to converge back to a single node per gene. For the simulations, the width of the network (the number of neurons in the layers) was chosen to be proportional to the input size. More specifically, the number of neurons, the basic computational units in a neural network, was set to be 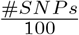 in the gene layer, with a minimum of five neurons. The learning rate for the ADAM optimizer and the strength of L1 penalty on the kernel weights were optimized on the validation set. To reduce the computational cost, hyperparameters were only optimized for a single simulation for simulations similar in sample size and input size. Networks were trained using CPU since the sparse matrix operations used do not benefit from using a GPU.

**Fig. 1.**
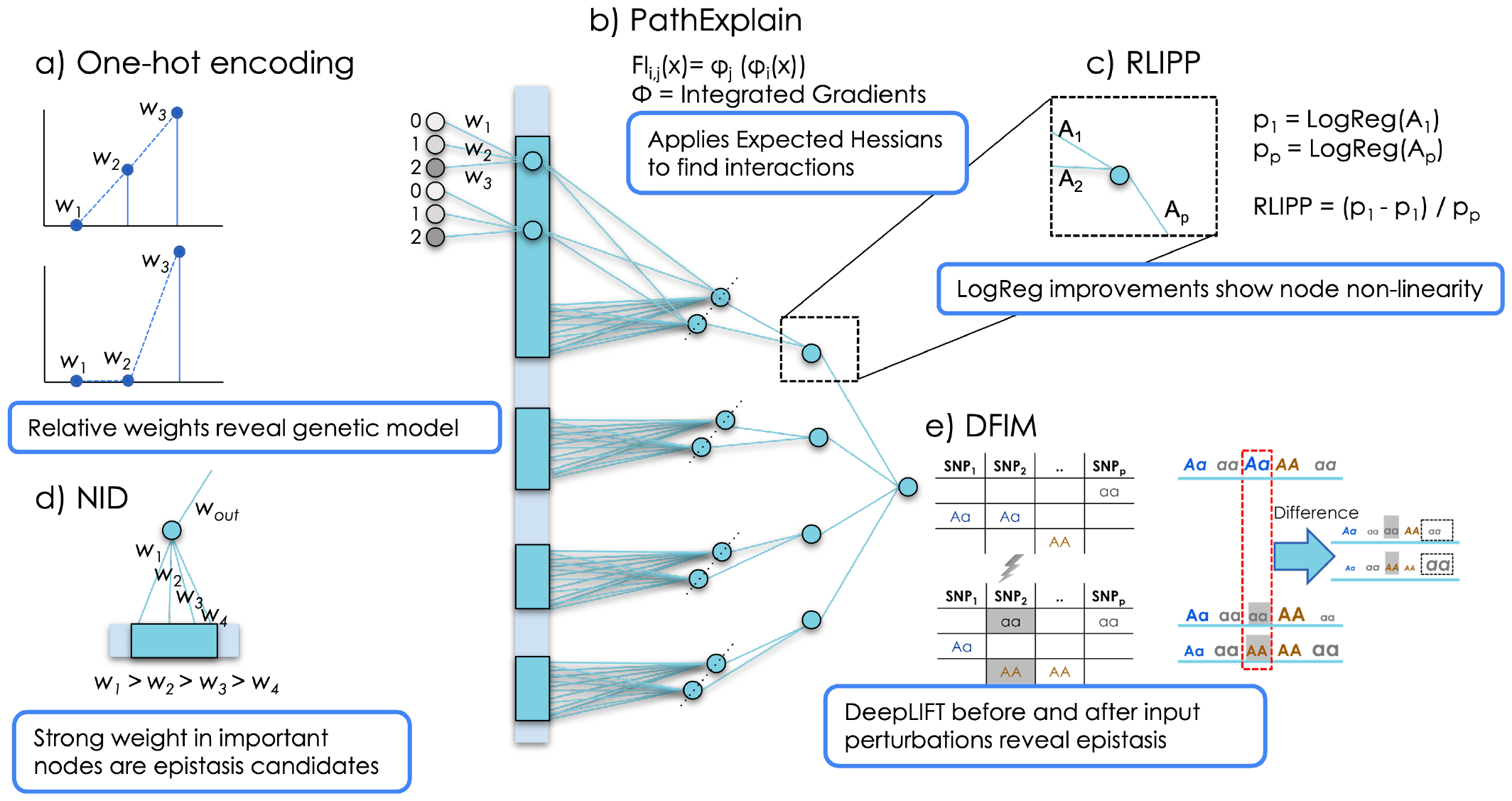
An overview of the post-hoc interpretation methods applied in this study to detect interactions in visible neural networks. (a) Comparing the relative weights of the one-hot encoded input for each SNP reveals the model that the neural network is using for that particular SNP (e.g linear spaced weights indicate an additive model). PathExplain applies Integrated Gradients on itself to find the Expected Hessians, which can be used to find interaction between inputs. RLIPP (c) is a method to detect if a node has non-linear behavior. The activations towards and from this neuron are regressed to the output with linear regression to provide an estimate of the non-linear gain of that node. (d) NID uses the assumption that edges with strong weights are more likely to interact with each other than edges with low absolute weights. DFIM (e) compares Deeplift’s attribution scores for all features before and after a feature of interest is perturbed, revealing all features that interact with the feature of interest.

For the IBD dataset we used similar neural networks but with gene annotations from FUMA (36). To map SNP to genes, both *positional* and *functional* annotations were combined. In the *positional* annotations, SNP to genes were mapped via a positional mapping obtained from FUMA’s SNP2GENE function. A SNP was mapped to a gene when the genomic coordinates of a variant were within the boundaries of a gene *±*10 kb. For the *functional* annotations, we used FUMA’s eQTL mapping that is based on eQTLs obtained from GTEx (37). An eQTL SNP was mapped to its target gene when the association p-value was significant in any tissue (FDR *<* 0.05). Combined they map 38225 SNPs to 25139 genes with 126899 connections.

### One-hot encoding

The standard genotype encoding {0,1,2} may introduce a bias to the additive model between genetic variants and the outcome as it represents an additive model. Therefore, we train for each application two models. A standard model and a model with a one-hot encoded input for the genotype. For this model we modify the first layer of each network, leading to three inputs per SNP (as shown in figure 1a). Regular one-hot encoding is widely used in machine learning to treat categorical variables as numerical values. In one-hot encoding the three categories that the SNP can assume: both reference alleles, a reference and an alternate allele, or both alternate alleles are each represented with a single variable that assumes value 1 if the input individual has said configuration, and 0 otherwise. We designed a novel way of using this one-hot encoded input, the three inputs per SNP are connected to a node representing that SNP in the first layer of the network. The weights of these connections may be informative of the genetic model learned by the model for that SNP.

For an additive model, the expected strength of the weights should roughly adhere to: *W*_0_ *−W*_1_ *≃W*_1_ *−W*_2_. We, therefore, use the ratio *R*_*w*_ as a measure for the degree of linearity. Historically, the additive model is the standard model, and it has had great success as the underlying model explaining genetic effects. Accordingly, we initialized the weights for each SNP according to the additive model, which can be seen as a reference model under Fisher’s epistasis definition. During training the neural network may freely change the ratio between these weights, diverging from an additive model. Inspecting the weights and the ratio *R*_*w*_ may indicate for which inputs the model deviates from the additive model. However, it is important to note that the model may learn more complex models using subsequent layers of the network, thus additive weights in the first layer do not exclude the possibility of a deviation of an additive model.

### Neural interaction detection (NID)

(27) is a method to detect statistical interaction pairs in neural networks that works on the premise that relevant interacting features have large weights assigned. Pairs of features are ranked according to the strength of the weights connecting to the neuron and the importance of the neuron (defined by the weights of its successive connections). Adapting this algorithm to visible neural networks is straightforward, as the mathematical interpretation is unchanged. The most likely interaction candidates are the combinations of the absolute weights that result in the highest value. Multiplying this value with the importance of the node, expressed by a multiplication of all the weights between the node and the output node, results in the final interaction score.

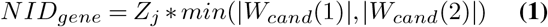

with *W*_*cand*_ as a matrix sorted by the absolute weights per gene and *Z*_*j*_ as the importance of the gene node (resulting from a multiplication of the absolute weights of all nodes between the selected node and the output).

### Deep Feature Interaction Maps (DFIM)

(26) assumes that perturbing a feature will result in a change in attribution score for a feature that is interacting with the perturbed feature. DFIM uses DeepLIFT (17) to get attribution scores before and after mutating a variant and saves this difference in attribution score as the feature interaction score (FIS). Since a single DeepLIFT call provides the attribution scores for all variants, only two calls are necessary to gain all the feature interaction scores for all the unperturbed (target) features. In this work, we perturb the hundred most important features identified by DeepLIFT and save the feature that has the highest FIS score, however a larger number of features can be saved if one suspects more interactions per feature.

### PathExplain

(28) uses the Expected Hessians for identifying interacting features. We apply PathExplain on the hundred most important features identified with expected gradient (38), the build-in feature importance method of PathExplain. For an input *x*, the feature interaction score (*FI*_*i,j*_(x)) is obtained using Integrated Gradients (*φ*) (16) applied on itself in order to explain the degree to which feature *i* impacts the importance of another feature *j*:

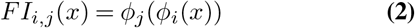

Thus, where DFIM mutates a feature and finds the change in importance for other features using the gradients in Deeplift, PathExplain directly finds the change in gradients by computing the expected Hessians.

### Relative local improvement in predictive power (RLIPP)

(13) is a method to detect in which nodes of the neural network statistical interactions occur. It compares the difference in predictive performance of a specific neuron’s inputs and outputs. The activations towards and from this neuron are regressed to the output with linear regression. In the original paper, the Spearman correlation was used to measure the performance gain for a regression task. We modified the algorithm in two ways to adjust it for the classification problem at hand. We compare the adjusted *R*^2^ of the two models and calculate RLIPP as:

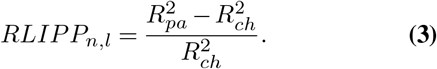

For GenNet’s networks for each node (*n*) and for each layer (*l*), the phenotype based on its’ activations (*Parent, pa*) and all the incoming edges, i.e., the weighted SNPs defining the node (*Child, ch*).

### Baseline methods

To compare the neural network to more traditional solutions, we use different epistasis detection models with different underlying mechanisms: LGBM, MB-MBDR and Epiblaster.

### Light gradient-boosting machine (LGBM)

(39) is a classification model using gradient boosting decision trees. Light gradient-boosting machine (LGBM) is an open-source gradient boosting framework that is designed to be efficient and scalable, making it well-suited for large datasets and high-dimensional problems. LGBM is particularly effective for handling large datasets with a large number of features, as it is able to handle missing data and categorical features efficiently. Two techniques distinguish LGBM from other gradient-boosting decision tree classifiers. *Gradient-based one-side sampling* allows LGBM to grow decision trees faster than traditional approaches, while also reducing overfitting and *exclusive feature bundling* to handle categorical features without the need for one-hot encoding, further reducing the memory usage and training time. We evaluated both feature importance as well as an unpublished feature interaction detection method (40), used in a Kaggle competition to predict customer transactions.

### Epiblaster

(21) is an algorithm that employs a two-stage approach to detect epistasis and generate a ranked list of SNPs with associated scores and adjusted p-values. The algorithm uses a combination of quasi-likelihood and linear models, such as linear regression or logistic regression depending on the output. In the first stage, an exhaustive filtering process is performed on all SNP pairs using the difference of Pearson’s correlation coefficient to rank them rapidly. In the second stage, only the top-k SNP pairs are selected, and a more accurate linear model with a real likelihood is used to compute the real p-value and test statistic. The p-value associated with the Beta is the measure of the association, and since multiple testing is performed, adjustment is needed. The retrieved p-value, adjusted for multiple testing, was used to rank the found SNP pairs.

### MB-MDR

(22) can identify genetic interactions in various SNP-SNP based epistasis models. The algorithm exhaustively explores the association between each SNP pair and the phenotype, using all available cases. It is a non-parametric method, in the sense that it makes no assumptions about the modes of interaction inheritance. The model-based part of MB-MDR refers to the ability to condition interaction testing on lower-order (main) SNP effects. For additional information and performance results, we refer to (41, 42); Such works report about the significance of interactions are corrected for multiple testing, using a step-down maxT inspired algorithm, controlling family-wise error rates (Type I errors). Exact permutation-based significance assessment is replaced by approximate such computations via the gammaMAXT algorithm as described in (43). By default, approximations are invoked when the number of input SNPs, to interrogate for interactions, exceeds 10, 000.

### Evaluation metrics

#### The Area Under the Receiver Operating Characteristic Curve (AUC-ROC)

is a popular metric used to evaluate the performance of binary classifiers. The ROC curve is created by plotting the true positive rate (TPR) against the false positive rate (FPR) for different thresholds. The AUC is simply the area under the ROC curve, ranging from 0 to 1, with an AUC of 0.5 representing a model equal to random guessing. We use the AUC to evaluate the classification performance of the neural network.

#### The Area Under the Precision-Recall Curve (prAUC)

is a useful metric to assess classifiers when there is a large imbalance between the classes. A high prAUC represents both high recall and high precision, where high precision relates to a low false positive rate, and high recall relates to a low false negative rate. The prAUC is used to evaluate epistasis detection.

### Simulation Results

We evaluated the performance and the consistency of interpretation methods for finding non-linear interactions with visible neural networks and compared these to more traditional approaches such as Epiblaster, MB-MDR and LGBM.

### GAMETES

The heritability value used in generating the simulations and the ease of detection, the difficulty based on the penetrance tables, had a clear impact on the predictive performance in the expected directions (see Supplementary Figure 5). To van Hilten and Melograna *et al*. | Detecting Genetic Interactions with Visible Neural Networks evaluate whether the post-hoc interpretation methods can extract the learned interactions in neural networks, we examined only simulations for which both types of neural networks found a predictive pattern, i.e., models with an AUC higher than an AUC of 0.5 in the test set (142/280).

Overall, the best interpretation method for the GenNet networks was DFIM with an average prAUC of 0.70 over all runs, followed by PathExplain (prAUC of 0.68) and NID (prAUC of 0.63) see (Figure 2a). There were strong correlations (Pearson correlation coefficients between 0.62 and 0.64) between the predictive performance of the neural network (AUC of the test set) and the ability of the interpretation networks to capture epistasis (e.g., DFIM prAUC). Figure 2b further dissects the relation between predictive performance and the ability of the interpretation methods to detect epistasis. In this figure, the performance of the various methods (NID, DFIM and PathExplain), are reported separately if they were trained with or without the one hot encoding. Moreover, it can be observed (Figure 2b) that for all networks with a prediction AUC of 0.6 or higher, DFIM and PathExplain achieved a prAUC of 0.98 and 0.95, respectively, with better results on the network without the one hot encoding. Interpreting the visible neural networks with NID resulted in a respectable prAUC of 0.89 for the same AUC threshold. There were negligible differences between standard GenNet networks and networks with a one-hot encoding in terms of classification performance. The networks with one-hot encoding did perform slightly better (average AUC of 0.64 vs 0.63) but the performance of the interpretation methods was worse, most noticeable for DFIM, where the average prAUC was 5 percent points lower for networks with the one hot encoding compared to networks that did not have the encoding. In addition, we performed simulations to investigate if these interpretation methods can find interaction between genes (gene-interaction). In these simulations the epistasis pairs are located in different genes. Classification performance was poor and dropped significantly compared to simulations where the interacting variants were in the same gene (see Supplementary Figure 6).

**Fig. 2.**
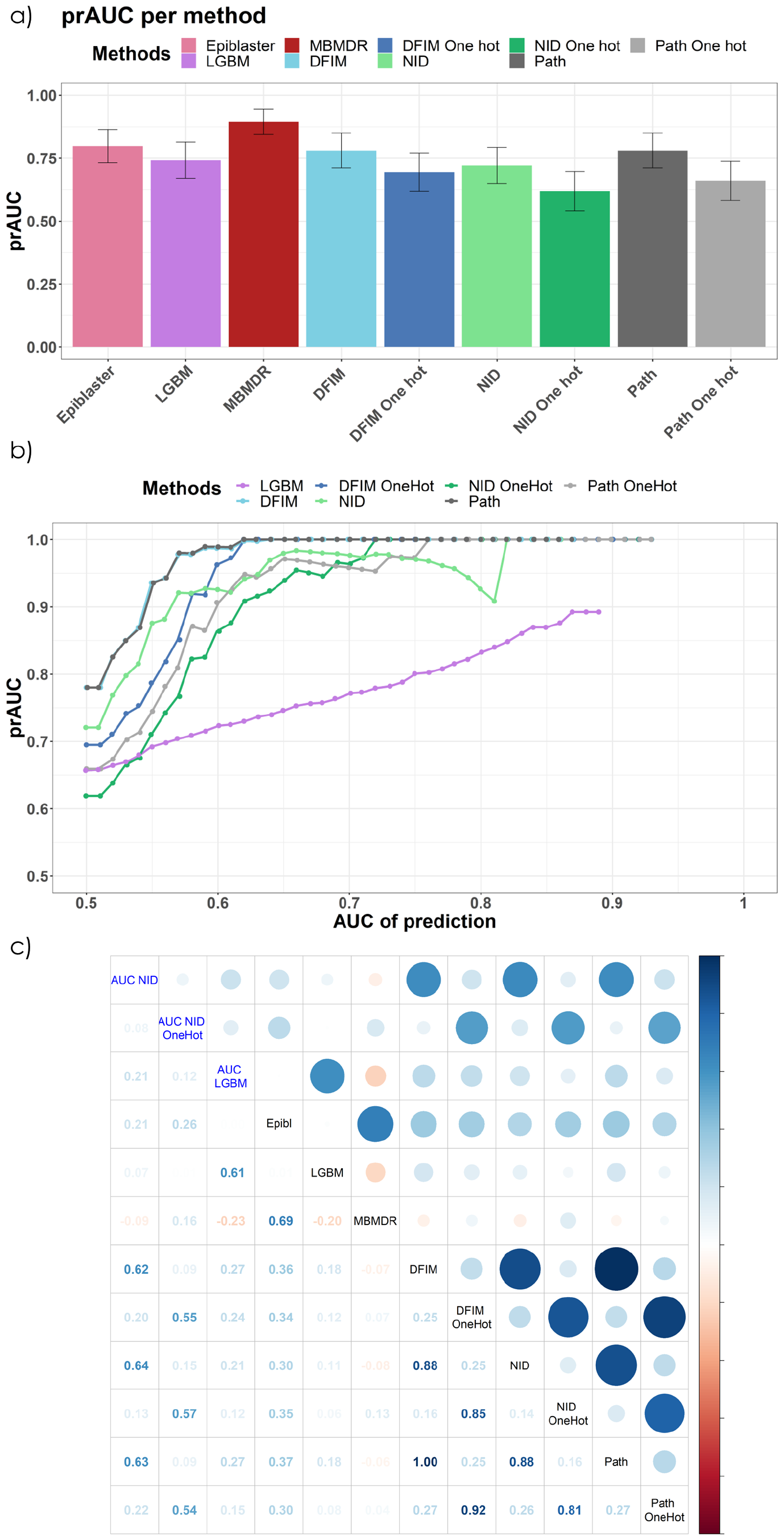
GAMETES: In a) the prAUC with the confidence interval of of the various epistasis interpretation methods.In b), the average of the prAUC for methods for different thresholds of prediction AUC in the test set. There is a clear trend showing better prAUC given better prediction AUC. In c), the correlation plot shows the correlation between the price of various methods and the prediction AUC of the NN and LGBM (AUC NN; AUC NN OneHot; AUC LGBM).

Baseline methods performed very well on GAMETES. LGBM slightly outperformed the neural networks in classification performance and for epistasis detection MBMDR outperformed the neural network interpretation methods in most simulations. Epiblaster achieved an average prAUC of 0.80 over all the simulations. Figure 2c displays the correlation between the results of these methods. We find a strong correlation (Pearson) between the predictive AUC of GenNet (blue), with the prAUC of the DFIM; NID and PathExplain. The same is true between the AUC of prediction AUC and the epistasis prAUC of LBGM.

### EpiGEN

The EpiGEN simulations were designed to investigate the behavior of the models for more realistic simulations with marginal effects and different interaction models. The interaction model strongly affects the classification performance of the GenNet models (see Supplementary Figure 7). Networks performed best in simulations using a *multiplicative* model, followed by *exponential* and *joint-dominant* models. *Joint-recessive* interaction models were the hardest types of interactions to capture in these sparse neural networks. In comparison to the GAMETES simulations, the performance difference between simulations with interacting pairs in the same gene versus interacting variants in different genes, was less pronounced, possibly due to the presence of marginal effects (Supplementary figure 7). The number of inputs and training-set size were clearly affecting predictive performance (Supplementary Figure 9).

To investigate the performance of the epistasis methods, we considered the subset of trained networks that achieved an AUC of 0.5 or higher for both types of networks. Figure **??** shows the average performance for each interpretation method. Predictive performance and interpretation performance were generally better for neural networks with a one-hot encoding than their corresponding networks without one hot encoding (see also Supplementary Figure 8 and 10). We found a similar positive trend as in the GAMETES simulation between prAUC and the AUC of the prediction 3c for all the neural network interpretation methods. However, thanks to the marginal effect, the neural networks can achieve a higher AUC with a poorer prAUC. Inspecting figure 3c shows that, for the same prediction AUC threshold, the prAUC is generally lower than in the GAMETES simulation (Figure 2c).

**Fig. 3.**
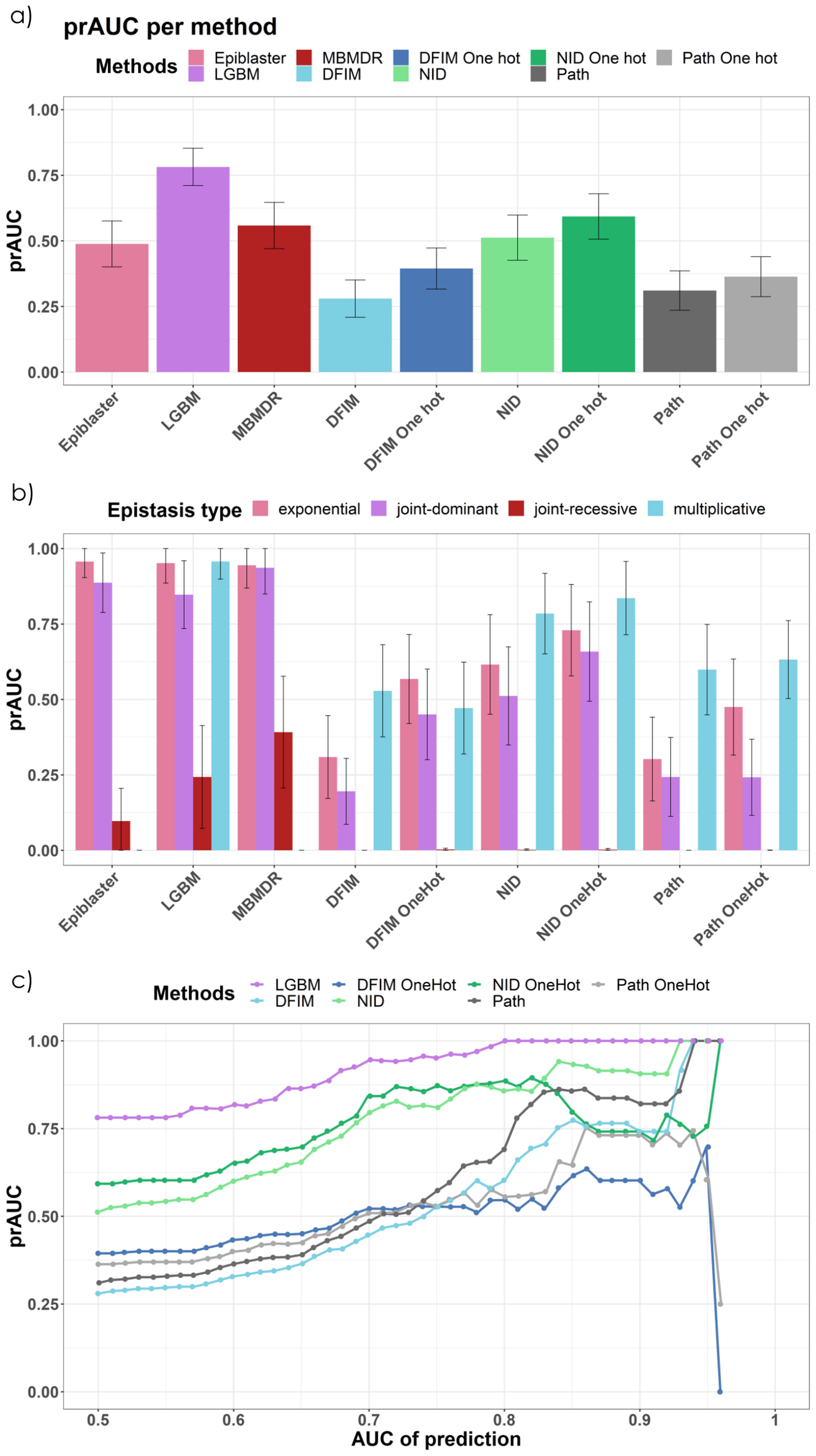
EPIGEN: In a), the mean prAUC of the various methods are compared, with the confidence interval displayed. In b), the mean prAUC of each method is displayed per type of interaction. In c), each dot is the average of the prAUC for methods that have a prediction AUC equal or greater than the number on the x-axis.

Inspecting the one-hot encoding reveals that the networks did encode the interaction models differently. Supplementary Figure 13 shows the deviations from linearity *R*_*w*_ per interaction model. The distribution deviates the most for the joint-dominant weights; causal *joint-dominant* pairs’ weights plateau for dosage input values 1 and 2, making them clearly separable from random or marginal weights (Supplementary Figure 14). Multiplicative and exponential weights were stronger for all inputs, but this was indistinguishable from the weight distribution for the one-hot encoding for variants with marginal effects. The weights distribution for the joint-recessive variants was most similar to those of random variants without any effects.

The best performing algorithm was LGBM; LGBM detects epistatic pairs with high prAUC in simulations with *exponential, joint-dominant*, and *multiplicative* interactions models 3b. All models struggle to detect epistatic pairs in simulations with an underlying joint-recessive model. For joint-recessive models LGBM is only second to MBMDR (MBMDR average prAUC for *joint-recessive*: 0.39). However, MBMDR and Epiblaster are unable to detect *multiplicative* pairs.

### Application to the IBD Dataset

To showcase the potential of our approaches in real-life data, we applied the methods to the IBD dataset with 66 280 observations and 38 825 SNPs after preprocessing. We divided the data into train (65%), validation (20%) and test (15%). Neural networks were created using GenNet command line functionality and both positional and functional annotations. As a result, a SNP can be linked to multiple genes. The covariates are inputs to the last hidden layer, before the final prediction. We built neural networks with and without one-hot encoding and achieved good predictive performance, with an AUC of 0.745 (0.715, for the one hot) in the validation set and 0.793 (0.761) in the test set.

### Interaction detection in visible neural networks

#### RLIPP

provides insight into which parts of the network the largest non-linearities can be found. We found that the node representing the gene *CCL11* had the highest relative improvement (see Supplementary Figure 11). *LYPLAL1-DT* and *SNX2P1* had the highest RLIPP values for the neural network trained with the one-hot embedding (see Supplementary Figure 12).

#### NID, DFIM, and PathExplain

The top epistasic pair (hit) of NID, for both the network built with and without the one hot encoding, was *rs2066844*-*rs2066845*, both missense variants in the *NOD2* gene and leading causal variants of Crohn’s disease and IBD in both Dis-GeNet and SNPedia databases (https://www.snpedia.com and https://www.disgenet.org). PathExplain, on the neural network with the one hot layer, had the SNP *rs*2066844 (NOD2) as part of the top epistasic pair together with *rs*5743293 (NOD2), a frameshift variant, related to both Chron’s disease (vda score = 0.83) and IBD (vda score 0.02). *rs*5743293 is particularly important for PathExplain one hot, as it is a hub involved in all the top-100 interactions. DFIM (on the network with one hot), showed the same behaviour, having a SNP, *rs*12946510, involved in 99 out of the top 100 interactions. *rs*12946510 (IKZF3, GRB7) is an intergenic variant associated to Crohn’s disease, IBD and Ulcerative colitis, as per the GWAS catalog. DFIM’s, on the one-hot neural network, top epistasic pairs involve *rs*12946510 (IKZF3, GRB7) with *rs*2066844 and *rs*2066845, both previously described. A list of the top SNP-SNP interactions for NID can be found in Supplementary Table 5 for the network without the one hot encoding layer and in Supplementary Table 4 for the network with the one-hot layer.

**Table 4.**
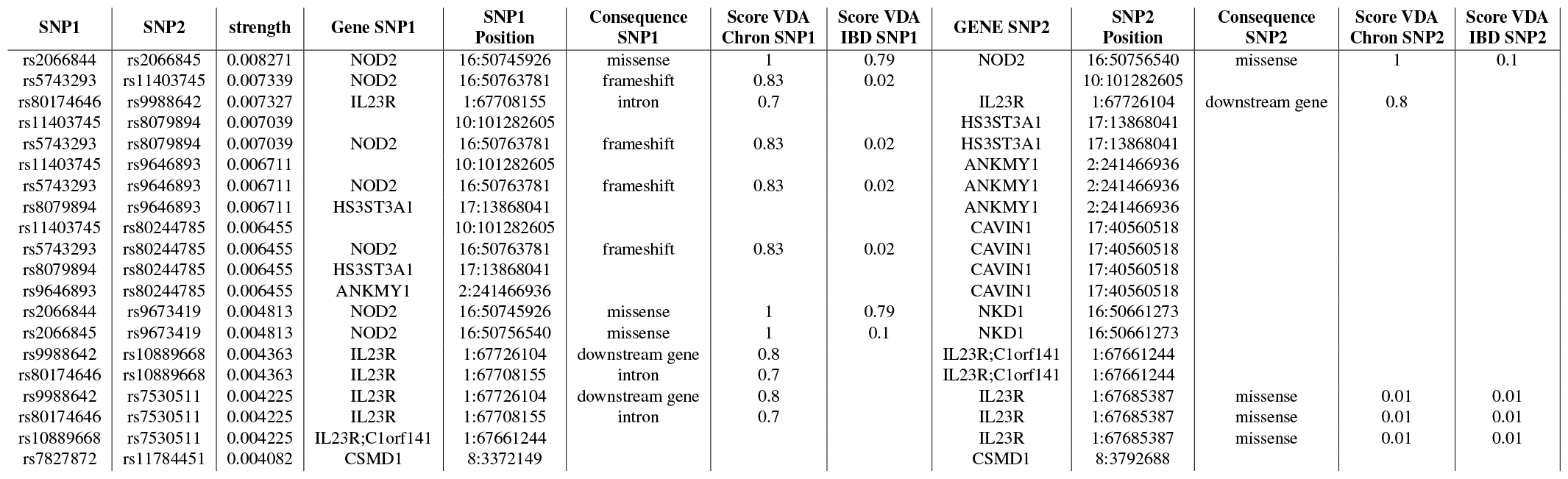
Top 20 SNP-SNP interaction for NID, trained with One-hot, as ranked per their strength. For each SNP in the pair, we mapped to the corresponding gene and, if available, we add the DisGeNet information on Chron’s disease and Alzheimer. In particular, we add the most severe consequence, the VDA score and the PMIDs, i.e., the number of studies were it was deemed relevant, the latter two for both Chron’s disease and IBD.

**Table 5.**
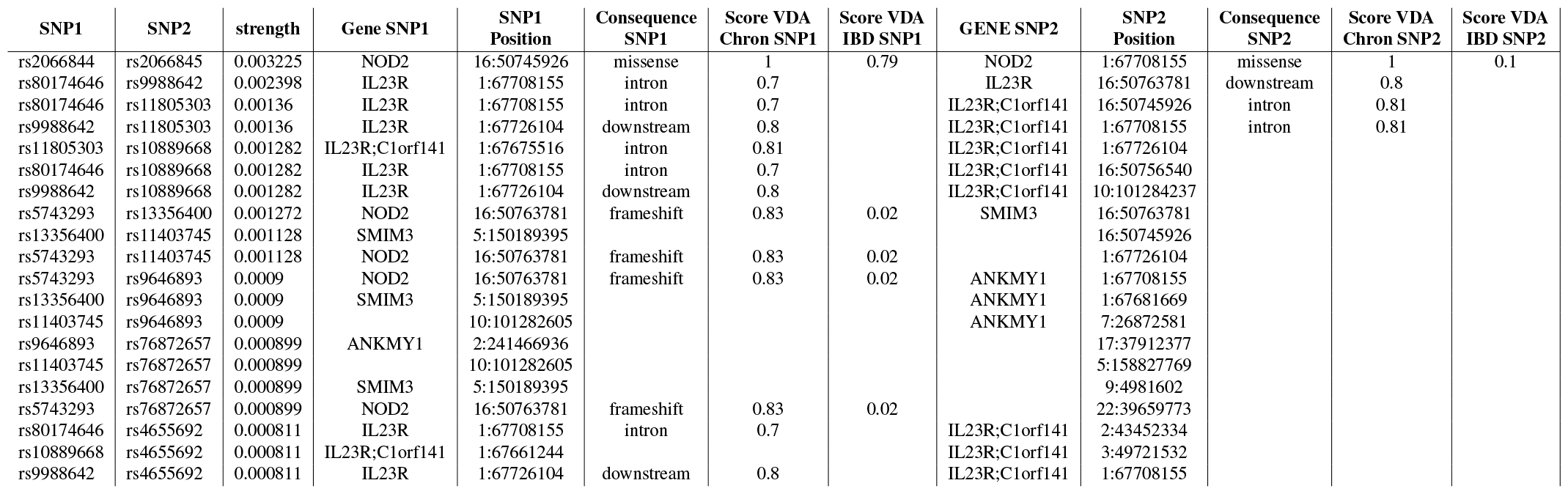
Top 20 SNP-SNP interaction for NID, trained without One-hot, as ranked per their strength. For each SNP in the pair, we mapped to the corresponding gene and, if available, we add the DisGeNet information on Chron’s disease and Alzheimer.

The top interaction in DFIM, in the network without the one-hot encoding, was between *rs*80174646 (intron variant, IL23R) and *rs*11805303 (intron variant, C1orf141/IL23R); both previously reported in association with Chron’s disease and IBD. (GWAS catalog). The second strongest interaction was between *rs*9988642 (IL23R) and *rs*11403745 (intergenic, LINC014675). The former is a downstream gene variant, mapped to the *IL23R* gene, a protein-coding gene associated with Inflammatory Bowel Disease. *rs11403745* is an intergenic variants whose closest gene is LINC01475, a non-coding gene. Nearby is also SEC31B, which has been associated to IBD. *rs*11403745 (intergenic, LINC014675) is also the SNP most present, 24 times, in the DFIM’s top 100 interactions. The same variant (*rs11403745*) is also part of the second top association in NID (on the network trained with the one-hot layer), together with *rs5743293* (NOD2), the hub SNP in PathExplain. Interestingly, a recent study highlighted *rs*11403745 in relation to IBD (44). *rs*9988642 (IL23R) and *rs*80174646 (intron variant, IL23R), part of the top and second interaction in DFIM (without one hot encoding layer), are also the second-highest interaction of NID without one hot encoding.

PathExplain on the network with the one-hot encoding detected the strongest interaction between *rs*9296009 (intergenic, closest are PRRT1, FKBPL) and *rs*2413583 (intergenic, RPL3, PDGFB). While the former has not been reported in the literature, *rs*2413583 has been associated with Chron’s disease, IBD, and ulcerative colitis, according to the GWAS catalog. Moreover, *rs*5743293 (NOD2) is the SNP most present, 26 times, in the top 100 interaction; it was part of all top 100 interactions of PathExplain with one hot layer and in the second position using NID on the network with the one-hot layer.

#### LGBM

There was no straightforward way to incorporate confounders into LGBM. Hence, we first regressed the phenotype with the 7 PCs with a linear model, subsequently using LGBM with the residuals as the outcome. LGBM provides both the feature importance and the interaction importance rankings for SNPs. Table 6 and 7 show the top-10 hits and the complete ranking can be found in Supplementary 6. Moreover, the most important feature according to feature importance, *rs2066844* (NOD2), is known to be the leading causal variant of Crohn’s disease. The top 3 features per LGBM’s feature importance, *rs*2066844, *rs*5743293, and *rs*2066845, are all linked to gene NOD2 and all associated with both IBD and Chron’s disease. Remarkably, out of the top-10 hits, 9 of them were already known in the literature to be associated with both Chron’s and IBD. The only hit not present in Dis-GeNet, *rs11403745* (intergenic, LINC014675) has been recently associated to IBD and has been extensively discussed in the previous subsection.

**Table 6.**
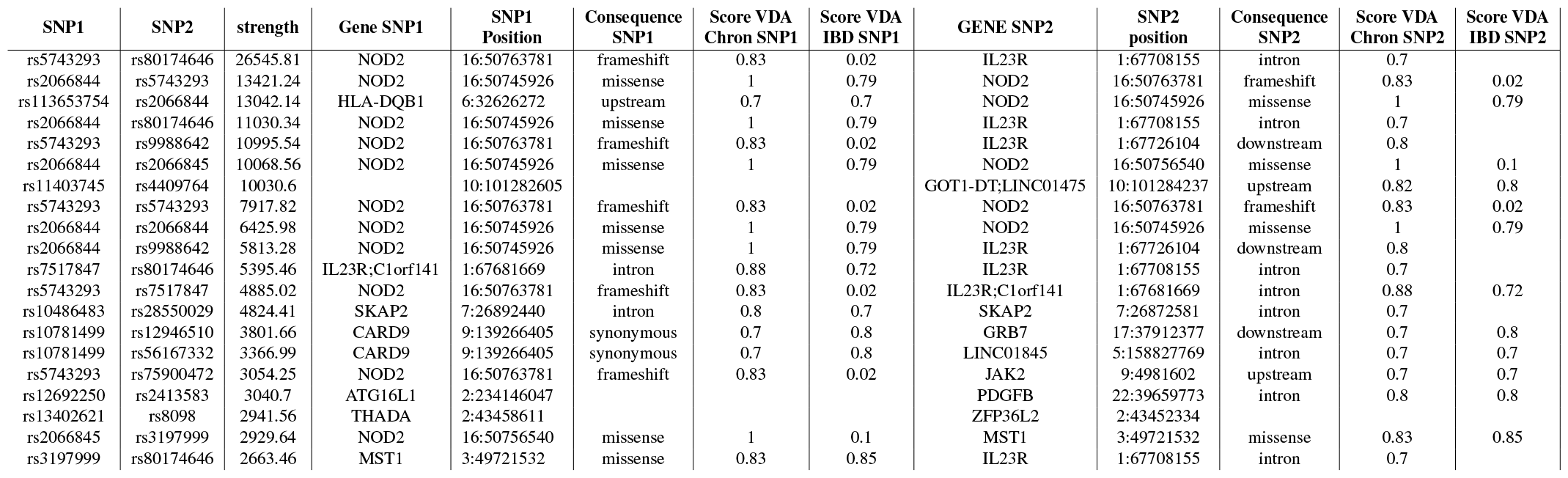
Top 20 SNP-SNP interaction for LGBM 2d, as ranked per their strength. It is worth noting that the strength for LGBM is calculated differently than NID and hence non-comparable. For each SNP in the pair, we mapped to the corresponding gene and, if available, we add the DisGeNet information on Chron’s disease and Alzheimer.

**Table 7.**
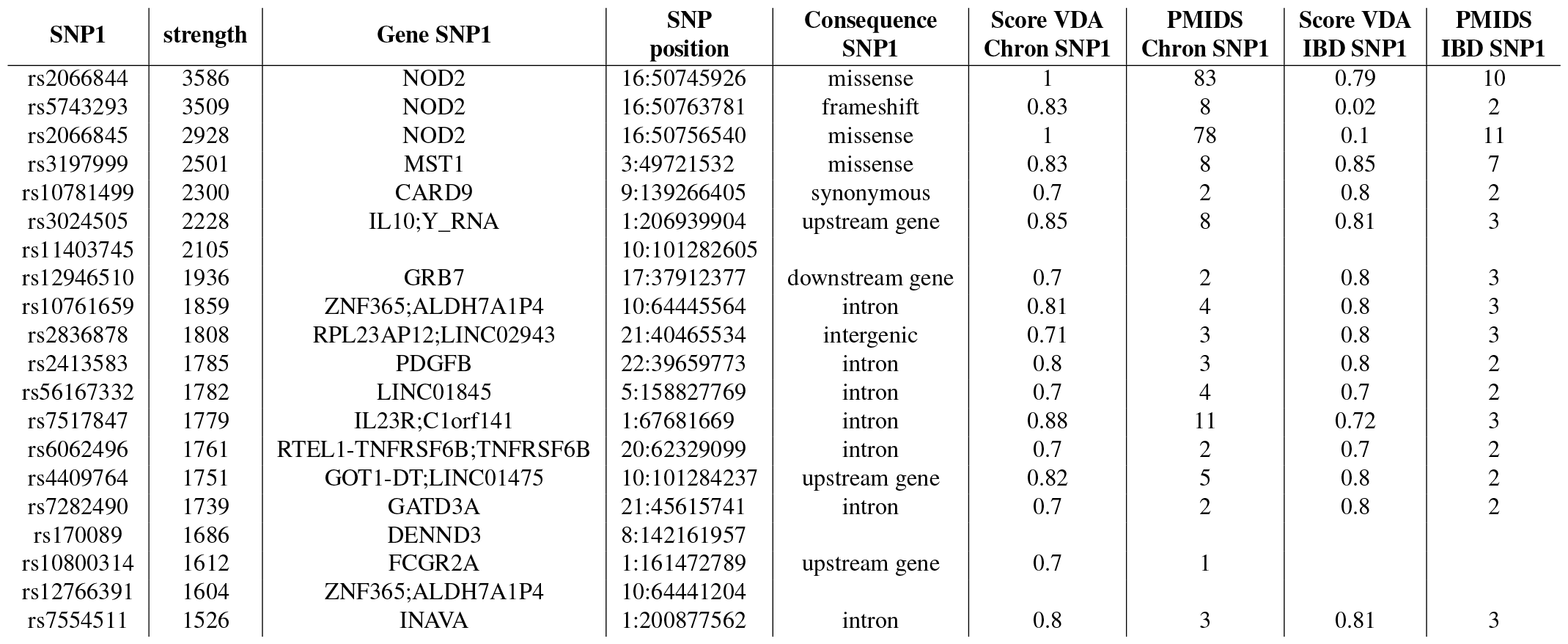
Top 20 SNP hits for LGBM 1d. We mapped each each SNP to the corresponding gene and, if available, we add the DisGeNet information on Chron’s disease and Alzheimer

For the LGBM’s interactions score, the top two SNP-SNP pairs involved *rs5743293* (NOD2), first with rs80174646 (IL23R); and then with *rs2066844* (NOD2). All three SNPs are known in the literature and have been found and described by the various neural network interpretation methods above. Overall, out of the top-10 SNP-SNP interactions, all but one SNPs are present in DisGeNet, for either IBD or Chron’s disease. The majority of them are mapped to either IL23R or NOD2. The single SNP that is not present in DisGeNet is *rs11403745* (intergenic, LINC014675).

### Rank aggregation and shared variants

Overall, we investigated the accordance and the peculiarities of each method on the IBD data for a broader picture of the agreement and disagreement of each interpretation method. We only calculated the interactions for the hundred most predictive variants for the DFIM and Pathfinder, restricting the search space, to reduce the computational burden.

First, we ranked every variant from NID, DFIM, PathExplain, both with and without one hot encoding layer, and LGBM, resulting in eight different rankings. For each method, the variant’s score is calculated as the sum of the interaction score (i.e., NID score, DFIM score, LGBM score,..) of every pair containing the variant. A comparative study from Li et al., (45) guides us toward using the geometric mean of the rankings. In this analysis, variants not present in a particular method, i.e., outside of the top-100 for DFIM and PathExplain, were assigned the lowest rank plus one. The geometric mean of the ranking of the eight methods highlights *rs*2066844, *rs*2066845, and *rs*5743293 (all NOD2 variants), as the top hits. Such variants were consistently present as top variants in each different method, with *rs*2066844 and *rs*2066845 ranked top-10 in 6*/*8 methods, with the only exceptions being 1) DFIM built without one hot layer and 2) the NID with the one hot layer.

Of the top-10 ranked hits, nine are already linked to either Chron’s disease (9) or IBD (7) in DisGeNet. The other hit is *rs*11403745, recently related to IBD (44). Another relevant SNP, in the top-100 in 7 out of 8 methods is *rs*9271588, a variant in the HLA region, that has been extensively studied in autoimmune disease and particularly Sjögren’s disease (46, 47). The full ranked table is available in the Supplementary Materials.

Furthermore, we plotted for each method the top 100 variants, the most promising candidates for epistasic effects, in an UpSet plot (Fig. 4a). To immediately visualize the adherence of our hits with the literature, we also included the list of SNPs associated with Chron’s and IBD present in DisGeNet that are part of our 38, 825 pool of input SNPs, respectively 314 and 228.

**Fig. 4.**
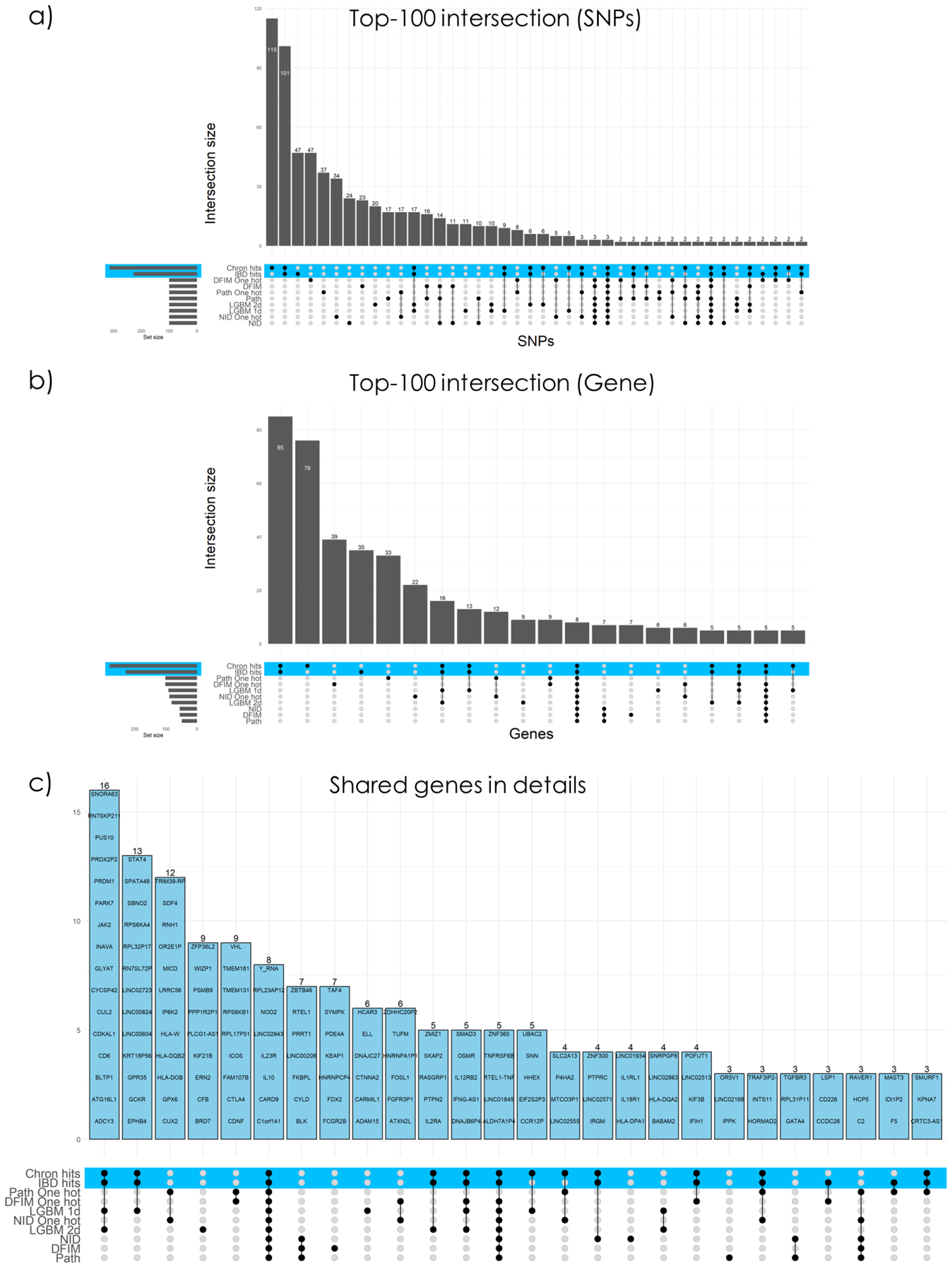
UpSet plot showing the intersections of our eight interpretation approaches (7 Epistasis methods: NID; DFIM; Pathfinder with/wihtout the one hot module, and LGBM’s feature interaction measure; plus LGBM feature importance) with the known variants from DisGeNet for IBD and Chron’s disease. Each standing bar shows the number of overlapping pairs between the highlighted method(s). In a), For each approach, the top-100 SNPs with the highest importance score were evaluated. The horizontal bar represents the number of SNPs included in each analysis, whereas the vertical bars show the overlap between each analysis; In b) the top-100 SNPs were mapped to gene positionally (as explained in the method section), and the intersection is showed. Finally, in c) the shared genes between at least one approach and one DisGeNet list are highlighted.

**Fig. 5.**
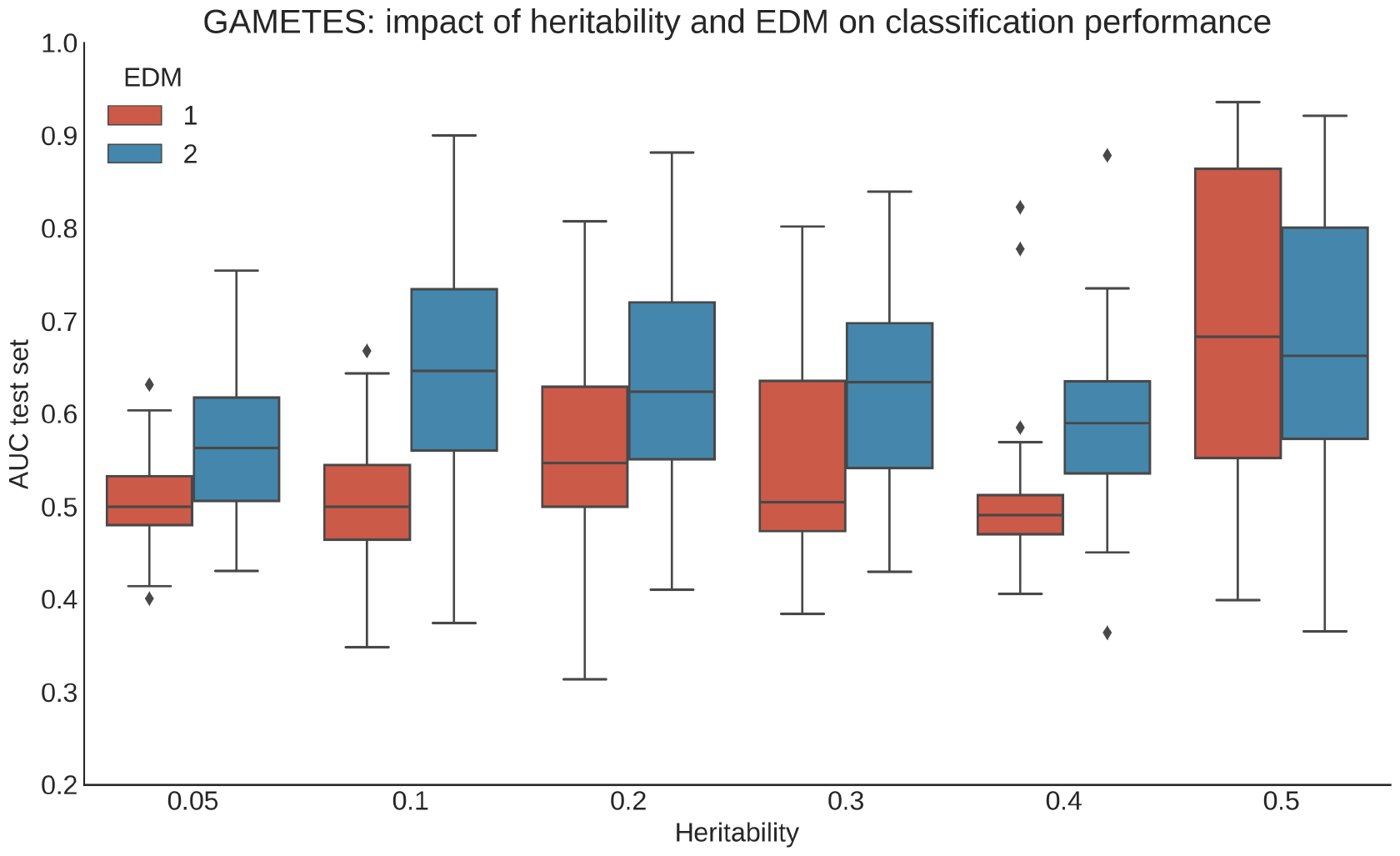
Boxplot of the predictive performance in area under the curve for the GenNet neural networks on the Gametes simulations for the different EDM and heritability parameters. EDM is a difficulty parameters of GAMETES.

**Fig. 6.**
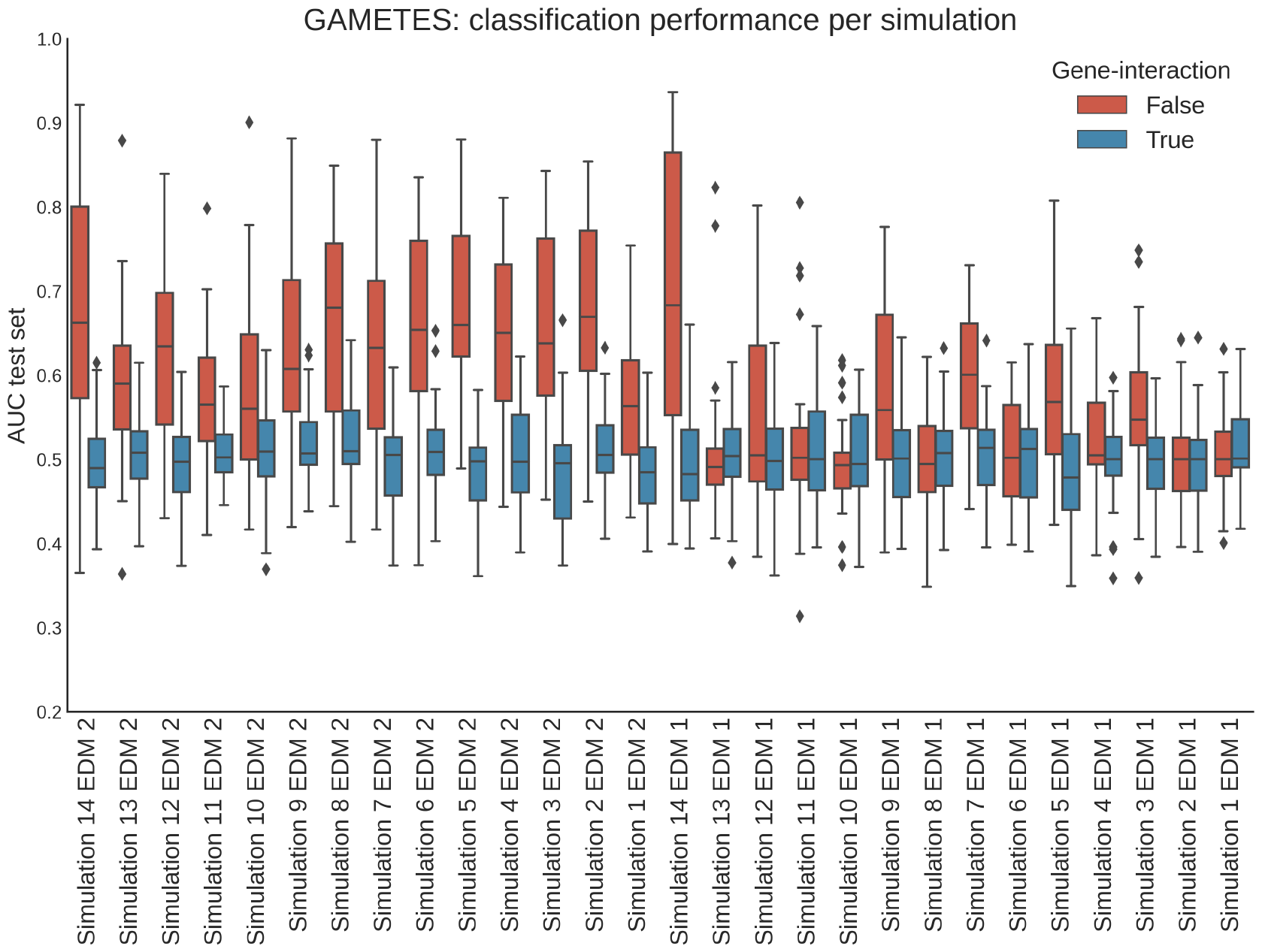
Boxplot of the predictive performance in area under the curve for the GenNet neural networks on the different Gametes simulations for the simulations with the interacting pair in the same gene and simulations with the interacting variants in a different gene (gene-interaction). See Supplementary Table 2 for the simulation specifications

**Fig. 7.**
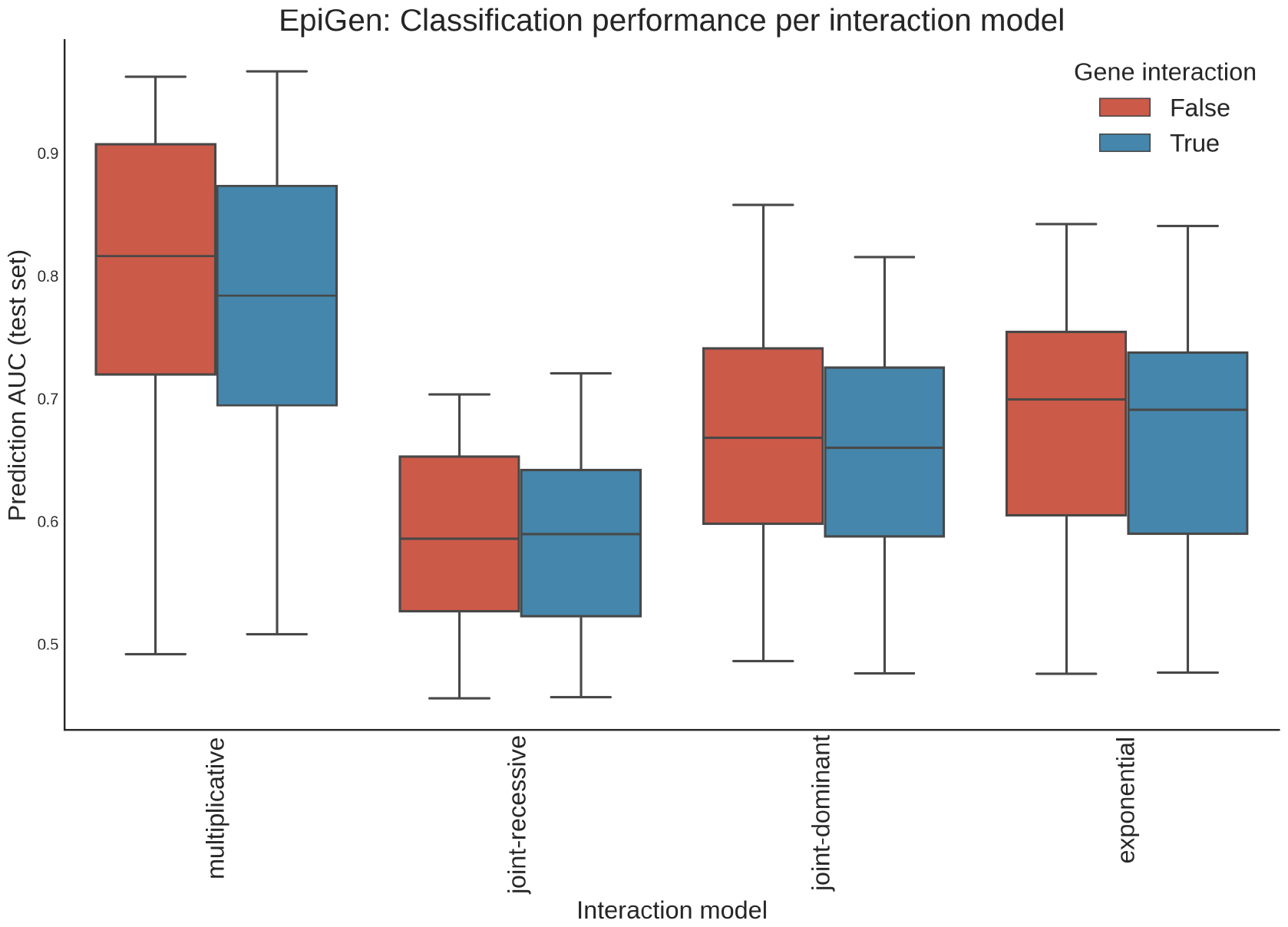
Predictive performance (AUC on the test set) of the GenNet models for various interaction models in EpiGen simulations colored by gene-interaction

**Fig. 8.**
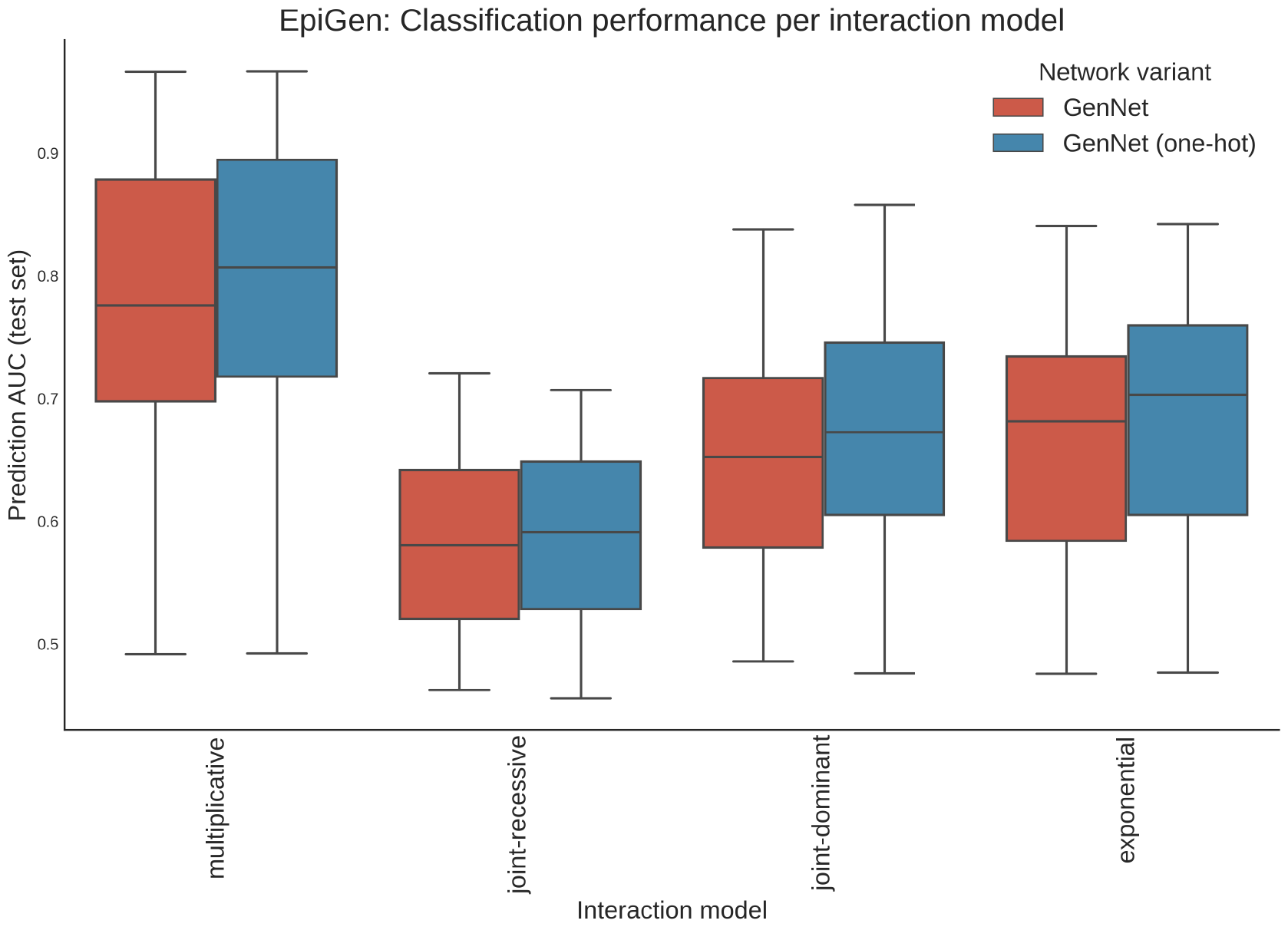
Predictive performance (AUC on the test set) of the GenNet models for various interaction models in EpiGen simulations colored by GenNet model. van Hilten and Melograna *et al*. | Detecting Genetic Interactions with Visible Neural Networks bioR*χ*iv | 15

**Fig. 9.**
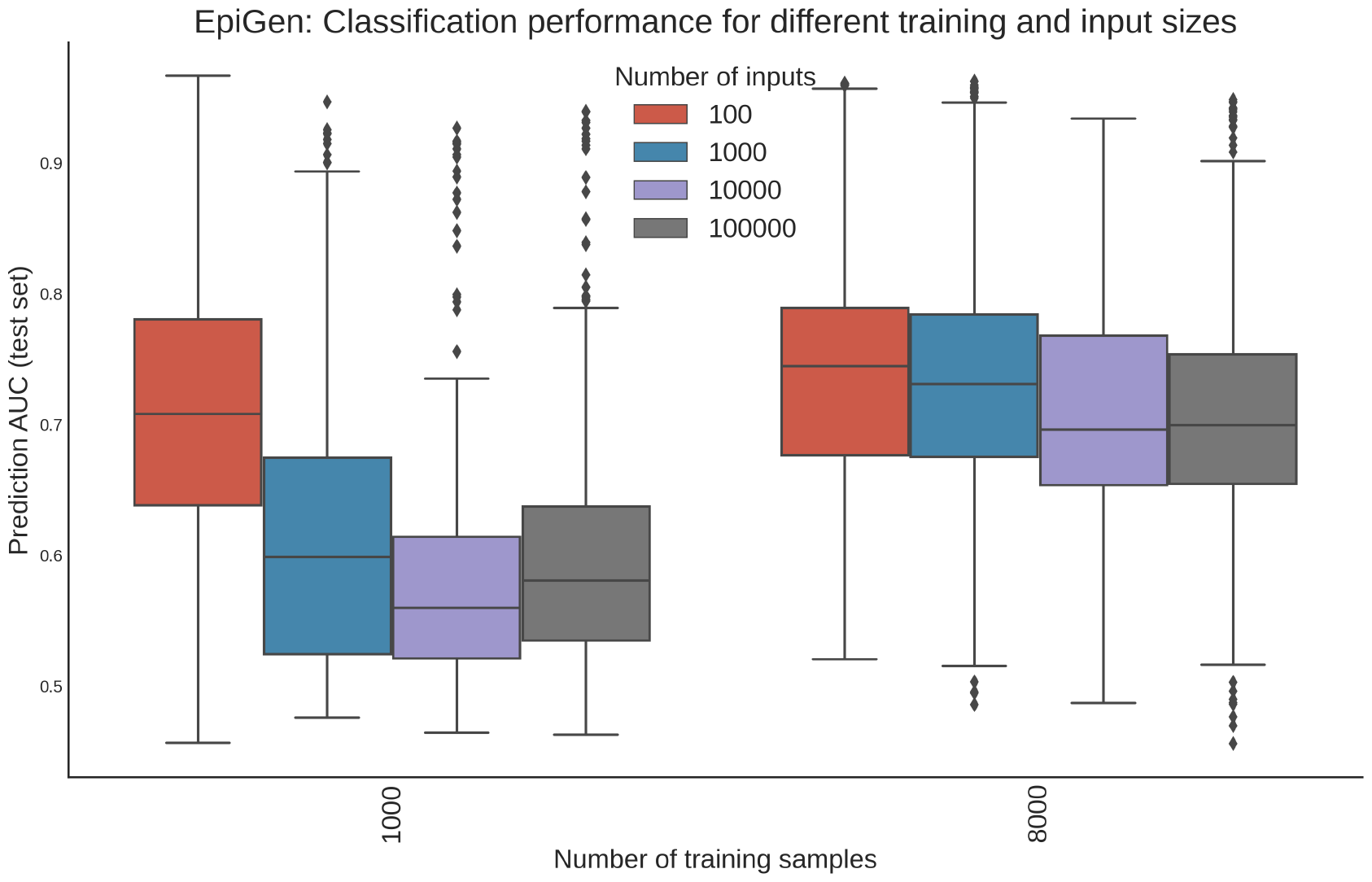
Predictive performance (AUC on the test set) of the GenNet models for various input and training sizes

**Fig. 10.**
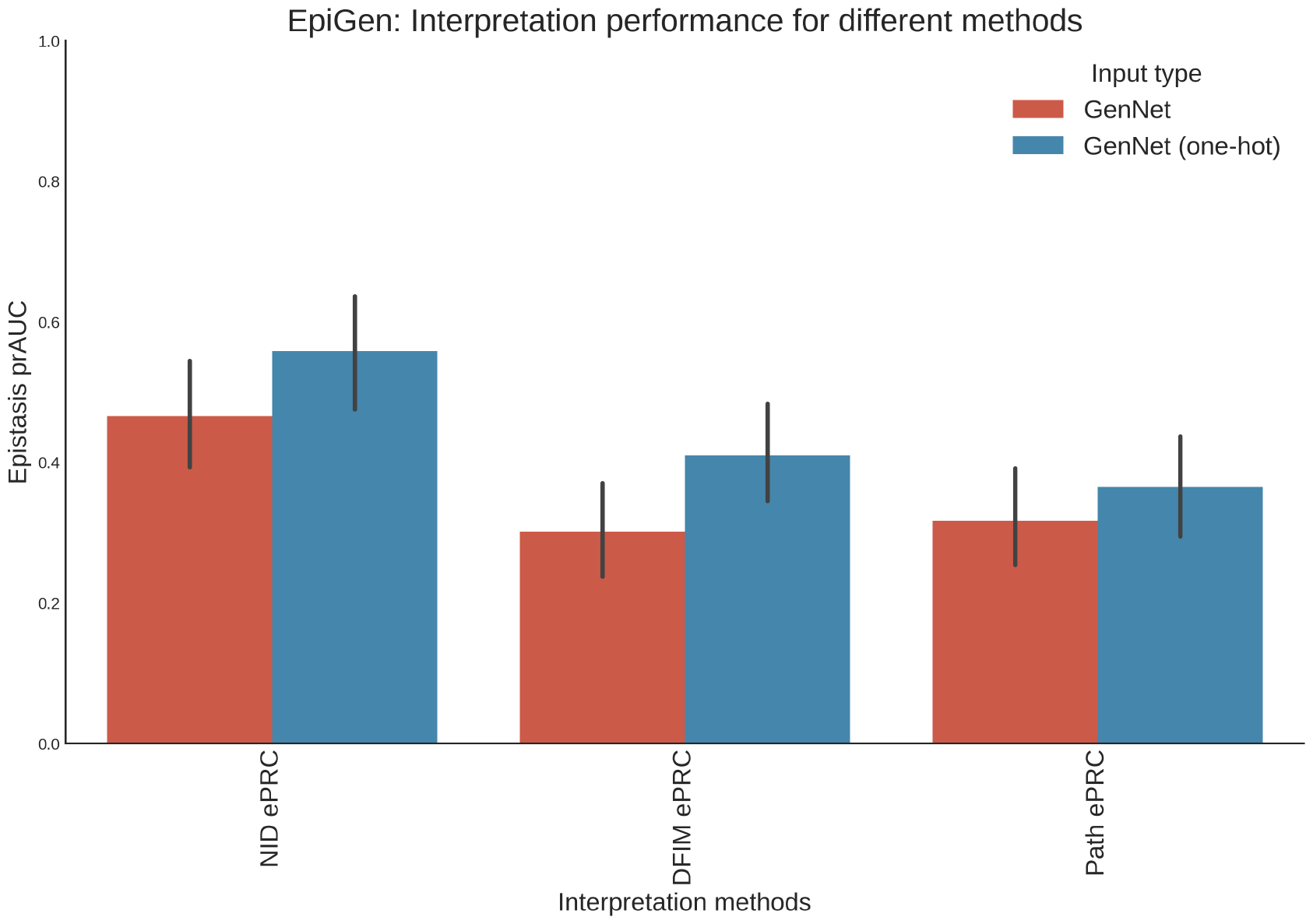
Predictive performance, AUC on the test set, of the GenNet models for various input and training sizes

**Fig. 11.**
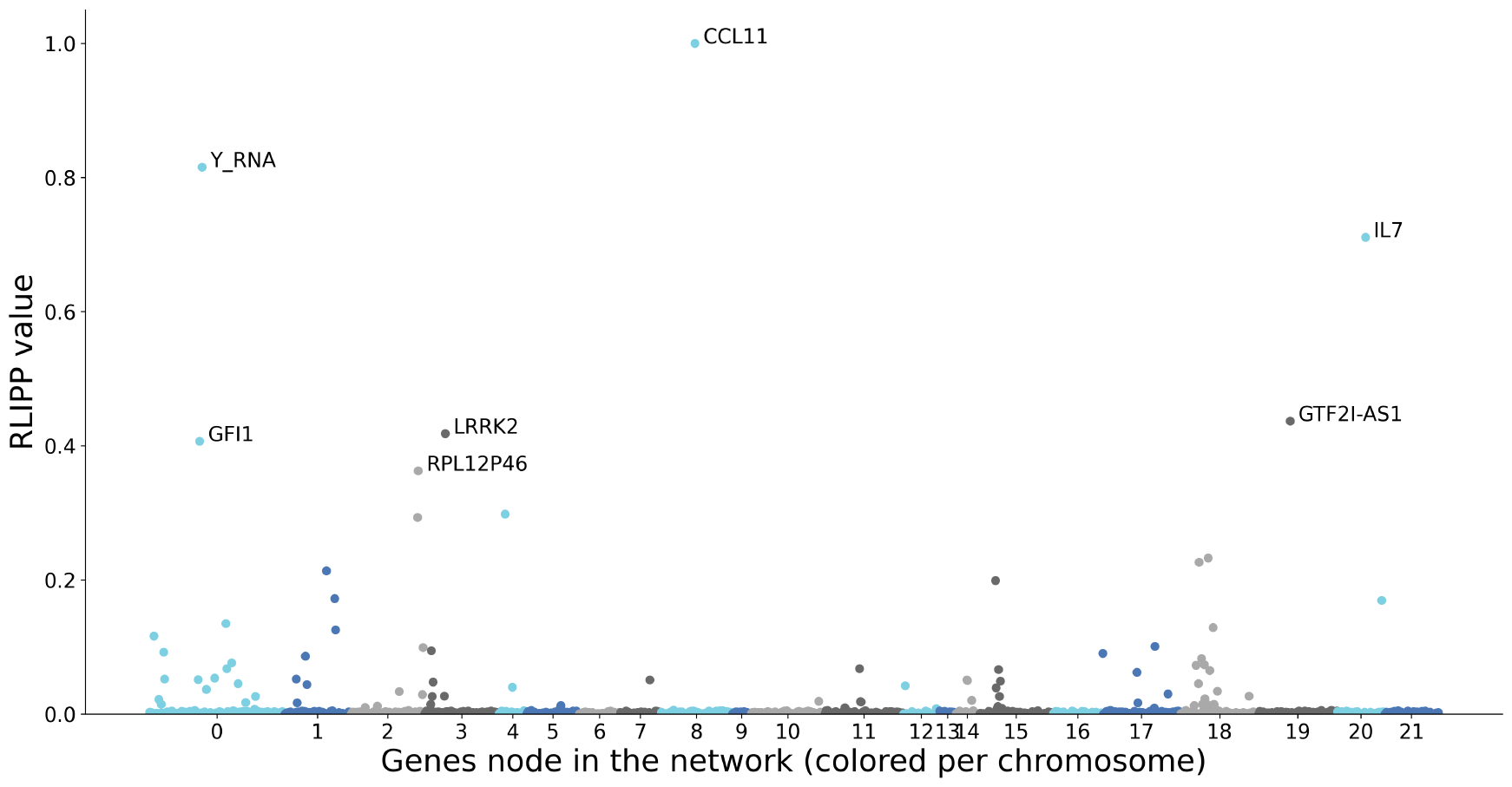
Normalized relative local improvement in predictive power (RLIPP) calculated for all gene nodes for the standard (additive) encoding of the neural network.

**Fig. 12.**
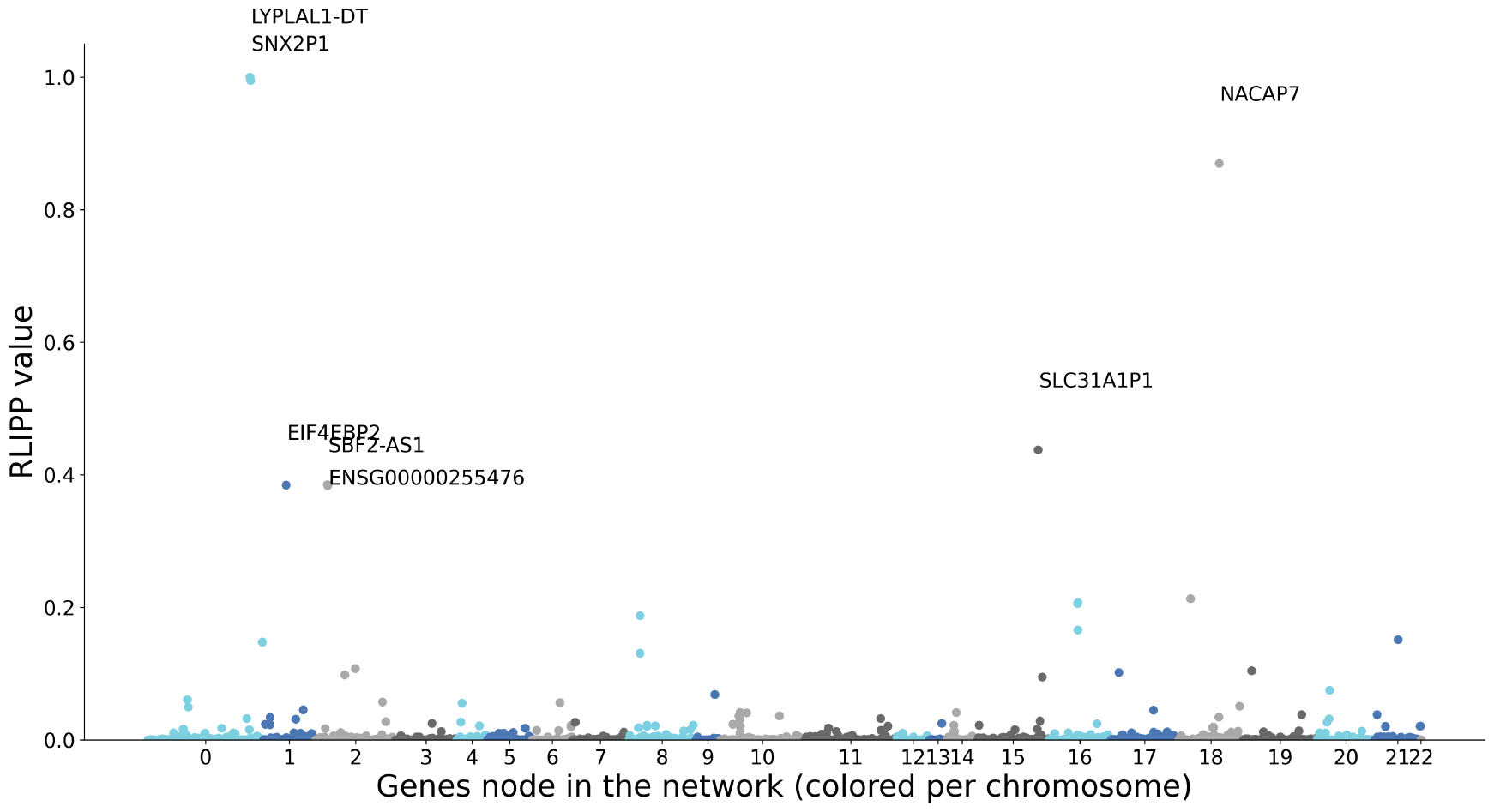
Normalized relative local improvement in predictive power (RLIPP) calculated for all gene nodes the neural network with the one-hot encoding. van Hilten and Melograna *et al*. | Detecting Genetic Interactions with Visible Neural Networks bioR*χ*iv | 17

**Fig. 13.**
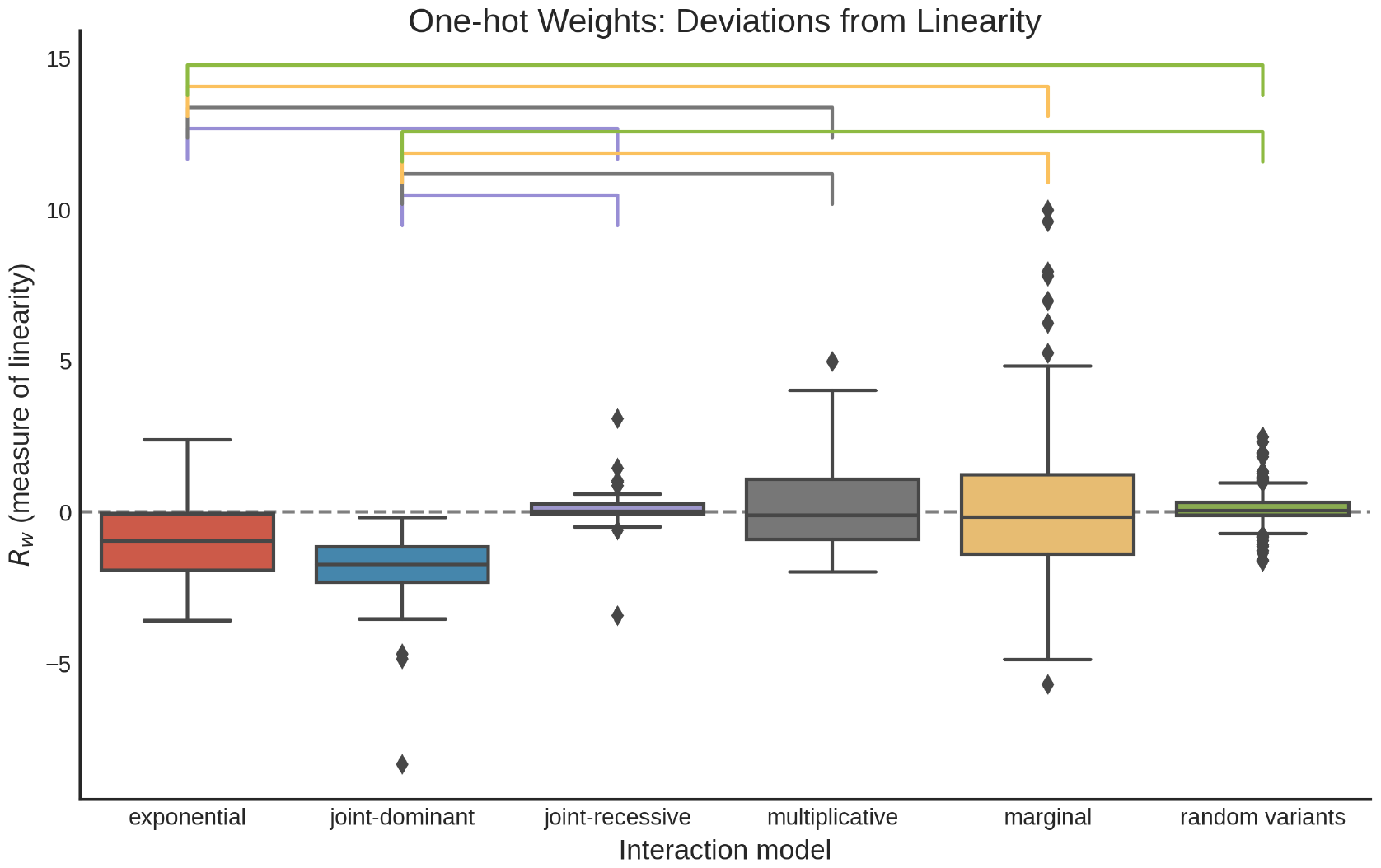
Boxplot with the deviations from linearity, calculated from the one-hot weights, for each interaction model. Significance is denoted on top (one way anova, p < 0.05)

**Fig. 14.**
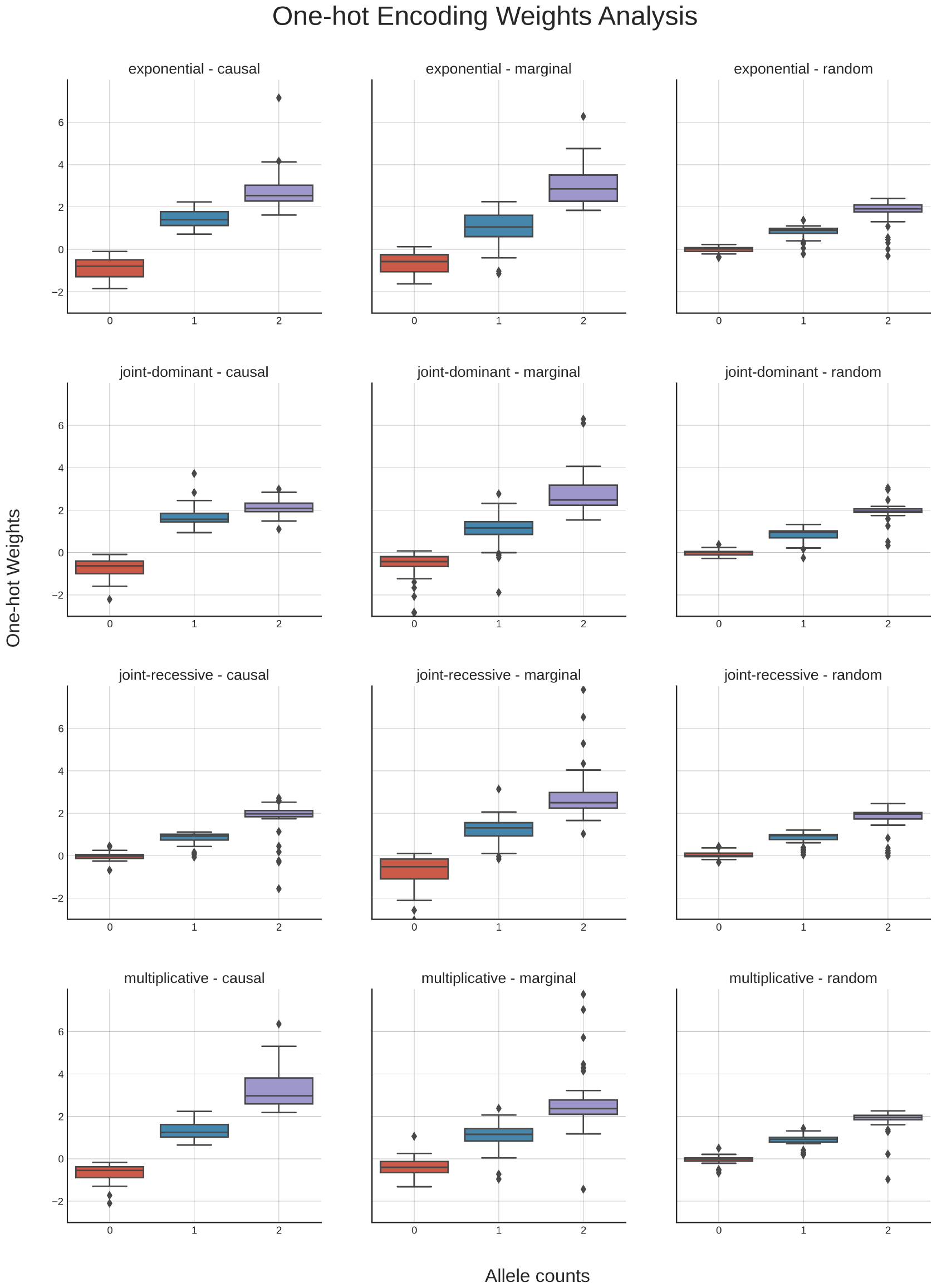
Weight distributions the one-hot encoding for the interaction models and all networks with an AUC > 0.6. Each columns show the causal variants, marginal variants and variants without effects, respectively. Each row shows the underlying interaction models for that simulation.

From this UpSet plot it can be seen that the top-100 hits combining all methods have a high overlap with the known hits in DisGeNet. LGBM’s feature importance (LGBM 1d) and epistasis detection (LGBM 2d) had the biggest overlap with *>* 40 out of the top-100 hits present in the DisGeNet hits for both Chron’s disease and IBD, respectively 62 and 55 for Chron’s and 52 and 45 for IBD. NID methods have around 20 hits, with, for NID NonOneHot and OneHot, respectively 23 and 24 in Chron’s disease and 17 and 21 in IBD. DFIM and Pathfinder have similar results, with the lowest number of hits belonging to DFIM NonOneHot on the IBD list, with only 12 hits.

Out of the considered methods, DFIM and PathExplain on the one-hot encoded network were the ones with the most unique hits, with DFIM having almost half of the variants in the top-100 not being in the top-100 of any other method or a known SNP from DisGeNet. On the other side of the spectrum, PathExplain on the NonOneHot and LGBM’s feature importance had the lowest number of unique hits.

Three variants were in the top-100 of all the mentioned methods, respectively *rs*2836878 (intergenic, RPL23AP12 and LINC02940), *rs*3024505 (upstream of IL10; close to Y_RNA), and *rs*10781499 (CARD9), with known association to IBD and Chron’s disease. The first is an intergenic variant, while the last is synonymous. Interestingly, in the GWAS catalog there are multiple studies linking *rs*10781499 to IBD disease, Ulcerative colitis and Chron’s disease. *rs*2836878 has also been associated with IBD, Ulcerative colitis, and Chron’s disease, as per the GWAS catalog. Finally, a study on a Danish cohort suggests a link between *rs*3024505 and the risk of Chron’s disease (48).

By mapping the top-100 SNPs to gene positionally (+/-10kb), we saw the overlap between methods and literature’s known hits (Fig. 4b). We found that eight relevant genes for both Chron’s and IBD (Y_RNA; RPL23AP12; NOD2; LINC02943; IL23R; IL10; CARD9; C1orf141) have at least one SNP mapped to them in each method (Fig. 4c).

### Association analysis for candidates pairs

We verified the findings from our previous methods with the most popular framework in epistasis detection, namely a logistic regression (LR). We grouped the top-100 SNP pairs from each of the seven epistasis methods. Hence, we ran a logistic regression to predict the phenotype using each pair of SNPs. The formula is, for a pair of SNPs *SNP*_*i*_ and *SNP*_*j*_, as follows:

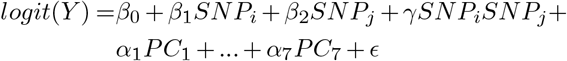

Where the *PCs* are the seven principal components to model population stratification. Hence, the *γ* coefficient reflects the epistasis interaction between a pair of SNPs. To avoid inflating the results, we ran logistic regression on the validation and test set combined, excluding the training examples that the network has seen.

Repeating the regression estimation for all pairs identified with the epistasis detection methods, we identified 7 significant SNP pairs after Bonferroni correction (Supplementary Table 8); out of those, two would stay significant under the usual GWAS threshold of 5 *∗* 10*−*8.

**Table 8.** Epistasis pairs found with logistic regression. In italics, the intergenic SNPs manually mapped to the flanking genes.

| SNP1 | SNP2 | Approach | pvalue | pos SNP1 | Gene SNP1 | pos SNP2 | Gene SNP2 |
| --- | --- | --- | --- | --- | --- | --- | --- |
| rs2294883 | rs454748 | LGBM | 4.93621E-09 | 6:32399674 | BTNL2;TSBP1-AS1 | 6:32245433 | *NOTCH4; TSBP1-AS* |
| rs11403745 | rs6584278 | LGBM | 3.40023E-08 | 10:99522848 | LINC014675 | 10:99518342 | *LINC01475; GOT1* |
| rs11403745 | rs4409764 | LGBM | 8.51701E-08 | 10:99522848 | LINC014675 | 10:99524480 | LINC01475 |
| rs2066844 | rs9673419 | NID OneHot | 1.1175E-07 | 16:50712015 | NOD2 | 16:50627362 | NKD1 |
| rs2066844 | rs5743293 | PATH OneHot,LGBM | 5.80966E-07 | 16:50712015 | NOD2 | 16:50729868 | NOD2 |
| rs1109863 | rs2066844 | NID OneHot, NID | 1.88448E-05 | 16:50658453 | *NKD1; LOC101927272* | 16:50712015 | NOD2 |
| rs5743293 | rs6715150 | PATH OneHot | 7.36191E-05 | 16:50729868 | NOD2 | 2:173890830 | SP3;LOC105373745 |

## Discussion

We adapted and applied various post-hoc interpretation methods to reveal the interactions learned by (visible) neural networks. Generally, we found that NID, DFIM and PathExplain are all suited to detect learned interactions from neural networks. There was a strong correlation between the predictive performance (AUC) and the ability of these interpretation methods to detect epistasis in the simulations (prAUC). That is, a neural network needed to have identified and learned the correct interactions before an interpretation method can extract it. There was no clear “best” interpretation method and the best interpretation method depends on the setting. In the GAMETES simulations, PathExplain performed best, while neural interaction detection (NID) was the best performing interpretation method for neural networks in most of the EpiGEN simulations. In the application to the inflammatory bowel disease, we found high agreement between the interaction interpretation methods. Interestingly, most variants identified by the interpretation methods were known variants earlier implicated in inflammatory bowel disease. From the candidate pairs identified with the interpretation methods on the neural networks and LGBM, 7 are significantly associated with IBD in the validation and test set.

In GAMETES, we empirically found that networks that achieved a classification AUC higher than 0.60 reliably detected interactions with most post-hoc analyses. Furthermore, the simulations revealed that interactions between variants located in different genes are hard to capture. The EpiGEN simulations confirmed both these findings and revealed that the ability to capture and detect epistasis pairs depends strongly on the underlying interaction model. Pairs based on a exponential model were consistently captured while pairs based on joint-recessive models were hard to model and detect. Increasing the depth of the neural networks, for example by adding pathway layers (10), may help with providing the networks with the necessary capacity to model interacting variants in different genes and with more complex interaction models.

There are large methodological differences between the methods employed in the simulations. The machine learning methods (neural networks and LGBM) optimize towards finding a good classification boundary, whereas MBMDR and Epiblaster are primarily designed to test for interaction effects. This could be an advantage for the simulations, as these methods align more closely with the process used to simulate the data and outcome. Both Epiblaster and MB-MDR can, however, be used in prediction models as part of a broader pipeline. For instance, prediction can be achieved via separating into training/test, 2) identifying (on the training set) the hits, both main effect and SNP-SNP pairs, 3) creating, for each observation, a weighted average of the hits’ effect; notable examples in the literature are (49), where MBMDR is used to build multilocus risk score (MRS) and MBMDRc (22), where the average trait for each SNP combination is averaged to build the prediction. It involves a generic strategy that could also be applied to other epistasis detection tools that do not readily provide predictions (such as Epiblaster). Hence, a notable difference is that in ML approaches prediction precedes the interpretation, while in epistasis tools it is the contrary.

In the IBD case-control setting, we achieved good predictive performance with both GenNet and LGBM. Interpretation revealed many variants that have been implicated to have a role in biological mechanisms underlying inflammatory bowel disease. This is likely a consequence of the initial filtering, narrowing the interaction interpretation down to pairs with at least one predictive SNPs in DFIM and PathExplain. NID did not require a filtering step as it is computationally cheap but the method inherently focuses on the variants with the highest weight. The most significant epistatic pairs are mapped to NOD2 variants (rs2066844, rs2066845,rs5743293) and IL23R variants (rs80174646). We confirm the recent finding of SNP rs11403745 (intergenic, LINC014675) for IBD, and propose variant rs9271588 (HLA region), as a candidate for further validation, being in the top-100 of 7*/*8 methods. Recently, Verplaetse et al. (50) applied biologically meaningful sparse neural networks on whole exome sequencing data to predict IBD. The authors achieved similar predictive performance but did not find convincing proofs for epistasis when comparing their performance to that of linear models. Here, we showed that by applying interpretation methods to the visible neural network we can detect epistasis. The reduced candidate set compensates with a lower multiple testing burden and thus more power, even-though half of the data is allocated for training the network and is thus unavailable for association analysis. Missing heritability is still a relevant problem for IBD and Zuk et al. (19) showed that up to 80% of the missing heritability could be due to genetic interactions. We detected 7 significant epistasis pairs in the real-life data but the simulations demonstrated that detecting epistasis pairs in different genes was difficult for the employed neural network architecture. Increasing the capacity of the neural networks to model these pairs could be a promising road for improving this strategy for epistasis detection.

We introduced several additions to the GenNet framework all of which, including the interpretation methods, are available from command line in the GenNet framework (https://github.com/ArnovanHilten/GenNet). We introduce multiple filters for visible neural networks, akin to channels in convolutional neural networks, and provide the option for an one-hot encoding for dosage input as a strategy to deal with the implicit bias to an additive model. With this encoding, the network is not forced to adhere to an additive model from the first layer and it is free to search for the encoding most suited for each single SNP. The one-hot encoding did result in minor performance gain in the EpiGen simulations and inspecting this layer revealed different weight distributions patters for the interaction models.

Here, we have demonstrated that interpretation methods for neural networks can identify non-linear interactions between genetic variants (epistasis pairs) in both simulated and real-life data. Most popular interpretation methods for neural networks provide a single importance (attribution) score per input, but this is inevitably a linear simplification of the true importance. Deep learning applications can model non-linear interactions and thereby provide a performance gain over linear models. In order to justify the use of these non-linear models it is thus necessary to use interpretation methods that can identify the non-linearities that lead to this performance gain. This does not only apply to epistasis; all tasks where neural networks are employed to leverage non-linear interactions can benefit from these interpretation methods.

## Conclusion

We demonstrated that interpretable neural networks can learn and detect epistasis using both simulated and real-life data. Moreover, we provided a comprehensive tool set and a novel strategy to interpret genetic interactions with visible neural networks.

## ACKNOWLEDGEMENTS

We would like to acknowledge all the investigators and participants in the International Inflammatory Bowel Disease Genetics Consortium. Funding was received from the European Union’s Horizon 2020 research and innovation programme under the Marie Sklodowska-Curie grant agreements N° 813533 (mlfpm.eu), N° 860895 (h2020transys.eu) and through the 2005 Simon Steven Meester grant 2015 to W.J. Niessen by the Dutch Technology Foundation (STW). Work was carried out on the Dutch national e-infrastructure with the support of SURF Cooperative (application number 17610). Gennady V. Roshchupkin supported by the ZonMw Veni grant (Veni 1936320)

## DATA AVAILABILITY

Code to run and generate data for the simulations are available on sourcefore (https://sourceforge.net/projects/gametes/files/) for GAMETES and on Github for Epigen (https://github.com/biomedbigdata/epigen) The genetic and phenotypic data for the IBD dataset are available upon application to the IBD consortium.

## CODE AVAILABILITY

All code for interaction detection (NID, RLIPP, PathExplain and DFIM) are now available in the GenNet interpret module. GenNet is an open-source framework usable from command line. GenNet can be found on: https://github.com/arnovanhilten/GenNet/ and Zenodo (). Epiblaster implementation used was https://github.com/FedericoMelograna/Epiblaster_implementation

## AUTHOR CONTRIBUTIONS

Arno van Hilten, Federico Melograna and Fan Bowen conceived, designed and performed the experiments. Gennady Roshchupkin, Kristel van Steen and Wiro Niessen supervised the work. Data set generation and quality control of the IBD dataset was done by the Inflammatory Bowel Disease Genetics Consortium. Details on contributions of all consortium members can be found on https://www.ibdgenetics.org/. Arno van Hilten and Federico Melograna wrote the first draft. All authors revised, and approved the paper.

**Supplementary Figures 1: GAMETES**

**Supplementary Figures 2: EpiGen**

**Supplementary Figures 3: RLIPP IBD dataset**

**Supplementary Figures 4: Epigen, one-hot encoding analysis**

**Supplementary Tables 5: IBD dataset**

**Supplementary Tables 6: Simulation characteristics GAMETES**

